# The landscape of nucleotide diversity in *Drosophila melanogaster* is shaped by mutation rate variation

**DOI:** 10.1101/2021.09.16.460667

**Authors:** Gustavo V. Barroso, Julien Y. Dutheil

## Abstract

What shapes the distribution of nucleotide diversity along the genome? Attempts to answer this question have sparked debate about the roles of neutral stochastic processes and natural selection in molecular evolution. However, the mechanisms of evolution do not act in isolation, and integrative models that simultaneously consider the influence of multiple factors on diversity are lacking; without them, confounding factors lurk in the estimates. Here we present a new statistical method that jointly infers the genomic landscapes of genealogies, recombination rates and mutation rates. In doing so, our model captures the effects of genetic drift, linked selection and local mutation rates on patterns of genomic variation. We then formalize a causal model of how these microevolutionary mechanisms interact, and cast it as a linear regression to estimate their individual contributions to levels of diversity along the genome. Our analyses reclaim the well-established signature of linked selection in *Drosophila melanogaster,* but we estimate that the mutation landscape is the major driver of the genome-wide distribution of diversity in this species. Furthermore, our simulation results suggest that in many evolutionary scenarios the mutation landscape will be a crucial factor shaping diversity, depending notably on the genomic window size. We argue that incorporating mutation rate variation into the null model of molecular evolution will lead to more realistic inferences in population genomics.

## Introduction

Understanding how various evolutionary mechanisms shape nucleotide diversity – typically measured as the average pairwise heterozygosity, π – is a major goal of population genomics (Charlesworth, 2010; Ellegren & Galtier, 2016), with a rich history of theoretical and empirical studies that have the fruit fly *Drosophila melanogaster* as its centerpiece (Casillas & Barbadilla, 2017; Charlesworth & Charlesworth, 2017; Haudry et al., 2020). For many years, the debate focused on the relative importance of genetic drift and natural selection to the genome-wide average π (Kimura, 1968; Ohta, 1992). The observation that π does not scale linearly with population size across species (Lewontin, 1974) was termed “Lewontin’s Paradox”, and recent work has taken a new stab at this old problem by modeling the effect of natural selection (Buffalo, 2021; Galtier & Rousselle, 2020). Later on, with recognition that linkage and recombination wrap the genome in regions of correlated evolutionary histories (Hudson, 1983; Hudson & Kaplan, 1985), focus shifted toward understanding how diversity levels vary along chromosomes of single species (Pouyet & Gilbert, 2021). In 1992, Begun and Aquadro found a positive correlation between π and local recombination rate in *D. melanogaster*, (Begun & Aquadro, 1992) which was interpreted as the signature of linked selection (Cutter & Payseur, 2013; Hudson & Kaplan, 1988) – at first in terms of selective sweeps (Smith & Haigh, 1974; Stephan et al., 1992; Wiehe & Stephan, 1993) and soon re-framed in the light of background selection (Charlesworth et al., 1993; Hudson & Kaplan, 1995, 1994; Nordborg et al., 1996). In the three decades since these seminal works, identifying the drivers of the genome-wide distribution of diversity became a leading quest in the field of population genetics. Nevertheless, this search has so far been incomplete. The literature has mostly considered how patterns of diversity are affected by selection (Andolfatto, 2007; Comeron, 2014; Elyashiv et al., 2016; McVicker et al., 2009; Murphy et al., 2022) or introgression (Hubisz et al., 2020; Stankowski et al., 2019), whereas spatial variation in *de novo* mutation rates (μ) has been largely ignored as an actual mechanism of variation in π along the genome, presumably due to challenges in its estimation (Besenbacher et al., 2019; Jónsson et al., 2018). Yet a study based on human trios advocates that the impact of the mutation landscape on polymorphism may be greater than previously recognized: up to 46% of the human-chimpanzee divergence, and up to 69% of within-human diversity, can potentially be explained by variation in *de novo* mutation rates at the 100 kb scale (Smith et al., 2018). It is unclear, however, how well these results generalize to species with distinct genomic features and life history traits. The few studies conducted in non-human organisms relied on proxies of the local mutation rate, such as synonymous diversity or divergence with a closely-related outgroup (Castellano et al., 2020, 2018). Still, these indirect measures of the mutation rate are susceptible to the confounding effect of selection, which can act both directly (e.g. codon usage (Lawrie et al., 2013; Machado et al., 2020)) and indirectly (e.g. recent background selection in the case of synonymous diversity (Charlesworth et al., 1993; Hudson & Kaplan, 1995; Nordborg et al., 1996) as well as background selection in the ancestral population in the case of synonymous divergence (Phung et al., 2016)). Therefore, developing dedicated statistical methods to infer mutation rate variation from polymorphism data is of high interest. Through simultaneous inference of the genomic landscapes of genetic drift, linked selection, recombination and mutation, confounding factors can be better teased apart and, subsequently, the relative contribution of each of these micro-evolutionary mechanisms to the distribution of diversity can be more meaningfully quantified.

Disentangling the effects of multiple factors shaping the evolution of DNA sequences is challenging because different mechanisms can produce similar phenomena (*sensu* (Baetu, 2019)). For example, a genomic region with reduced nucleotide diversity (relative to some baseline reference) can be causally explained by either linked selection, drift, low mutation rate or a combination thereof. In an elegant effort to tease these mechanisms apart, Zeng and Jackson developed a likelihood-based model that jointly infers the effective population size (*N*_e_) (Charlesworth, 2009) and μ with high accuracy in different parts of the genome (Zeng & Jackson, 2018). However, since it relies on the single-site frequency spectrum, their method is restricted to unlinked loci. While this approach avoids the confounding effect of linkage disequilibrium in the inference procedure (Slatkin, 2008), it discards sites in the genome where local variation in the mutation rate may be relevant as well as dismisses the gradual impact of recombination and linked selection on spatial variation in diversity. In this article, we put forward a new model to fill in this gap. We have previously described a statistical framework (the integrative sequentially Markovian coalescent, iSMC) that jointly infers the demographic history of the sampled population together with variation in the recombination rate along the genome via a Markov-modulated Markov process (Barroso et al., 2019). We now extend this framework to also account for sequential changes in the mutation rate. This integration allows statistical inference of variation along the genome in both recombination and mutation rates, as well as in Times to the Most Recent Common Ancestor (τ), that is, the ancestral recombination graph of two haploids (Rosenberg & Nordborg, 2002). Whereas drift causes stochastic fluctuations in τ around its expected value under neutrality (in diploid organisms, 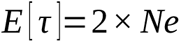), natural selection disturbs τ away from its neutral distribution near functionally constrained regions of the genome (Palamara et al., 2018; Rasmussen et al., 2014; Stern et al., 2019; Zeng & Charlesworth, 2011). Thus, iSMC offers estimators of all relevant micro-evolutionary mechanisms, and we can further use causal inference (Pearl & Mackenzie, 2018) to simultaneously estimate their effects on diversity. Our analyses of *D. melanogaster* genomes reveals the impact of linked selection; however, it suggests that the rate of *de novo* mutations is quantitatively the most important factor shaping nucleotide diversity in this species.

## Methods

### Modeling spatial variation in θ

We now introduce our approach to modeling the mutation landscape starting from the original pairwise SMC process. Because iSMC models pairs of genomes, the genealogies underlying each orthologous site can be conveniently summarized by τ, the time to their most recent common ancestor (Li & Durbin, 2011; Schiffels & Wang, 2020). The pair of DNA sequences is described as a binary string where 0 represents homozygous states and 1 represents heterozygous states (thus, once genome pairs are combined into diploids, phasing information is discarded). The probability of observing 0 or 1 at any given position of the genome depends only on τ and the scaled mutation rate θ. If the hidden state configuration of the model excludes spatial variation in the mutation rate, then θ is assumed to be a global parameter such that the emission probabilities of homozygous and heterozygous states can be computed for every site as 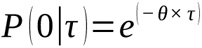, and 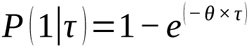 respectively, as presented by Li & Durbin (2011).

We estimate the per-site, genome-wide average 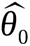 as the average number of pair-wise differences observed between all pairs of genomes. Therefore, the effective population size implicit in 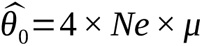 is the average of *N*_e_ along the genome, accounting for selective effects. We fix 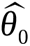 to this point estimate and exclude it from the optimization step conducted with the HMM. To incorporate spatial heterogeneity in the mutation rate along the genome, we modulate 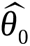 by drawing scaling factors from a discretized Gamma distribution with mean equal to 1. The parameter shaping this prior distribution (α_θ_ = β_θ_) is estimated by maximum likelihood (via the forward HMM algorithm) together with other parameters of the model (using the Powell optimization procedure (Powell, 1964)). We model the changes in mutation rate along the genome as a Markov process with a single parameter δ_θ_, the transition probability between any class of mutation rate, which is independent of the genealogical process. The justification for the Markov model is that sites in close proximity are expected to have similar mutation rates. For example, as is the case when the efficiency of the replication machinery decreases with increasing distance from the start of the replication fork (Francioli et al., 2015). Of note, Felsenstein & Churchill (1996) used a similar approach to model substitution rate variation across sites in a phylogenetic model. Let 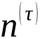 be the number of discretized τ intervals, and 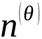 be the number of discretized categories of the prior distribution of scaling factors of θ. The ensuing Markov-modulated HMM has 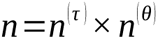 hidden states. The transition matrix for spatial variation in θ is:

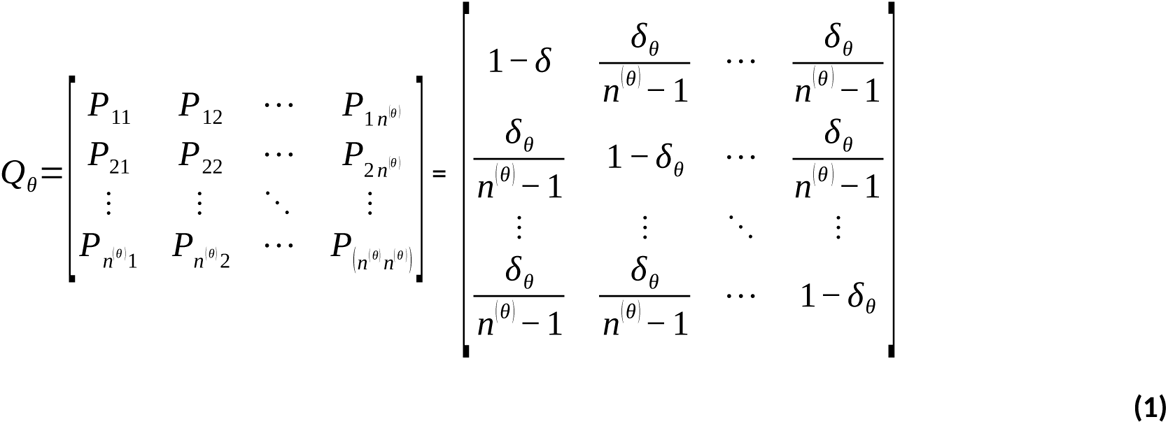

where δ_θ_ is the aforementioned auto-correlation parameter. The resulting process is a combination of the SMC and the mutation Markov model, so that its transition probabilities are functions of the parameters from both processes, that is, the coalescence rates (parameterized by splines, similarly to (Terhorst et al., 2017)), δ_θ_ and the global recombination rate ρ (Barroso et al., 2019). The forward recursion for this model evaluated at genomic position *i* can be spelled out as:

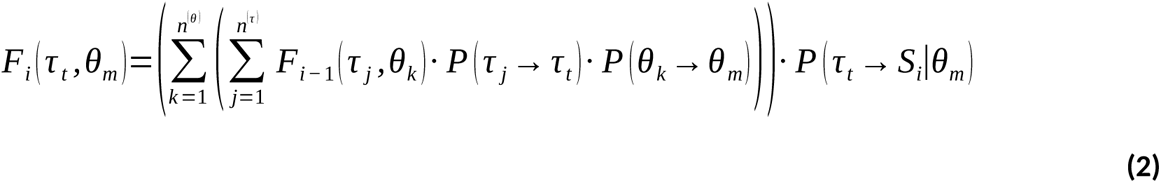

where θ_m_ is the product of 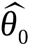 and the value of the *m*-th discretized category drawn from its prior Gamma distribution. The emission probability of binary state S_i_ depends on the height of the *t*-th genealogy and the focal mutation rate θ_m_. More specifically, the emission probabilities of θ-iSMC are 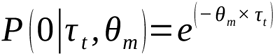, and 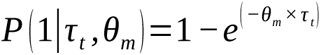. Thus, the forward recursion integrates over all 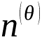 categories of θ and over all 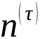 intervals of τ, for all sites in the genome. In the double-modulated model (ρ-θ-iSMC), where both mutation and recombination are allowed to vary along the genome, this integration is performed over θ, τ as well as ρ (giving a total of 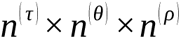 hidden states, **Figure 1**). Since spatial variation in ρ contributes to the transition probability between genealogies, the complete forward recursion is now given by:

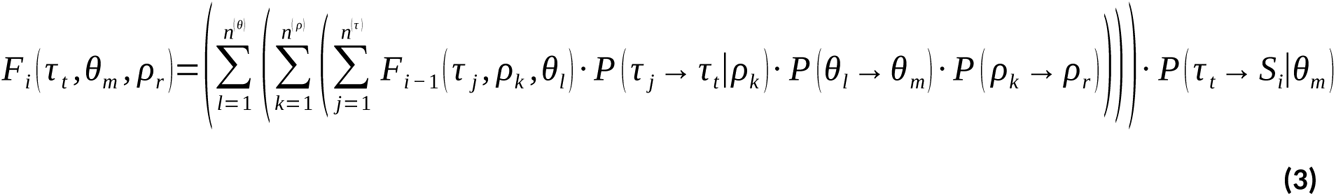

**Figure 1.**
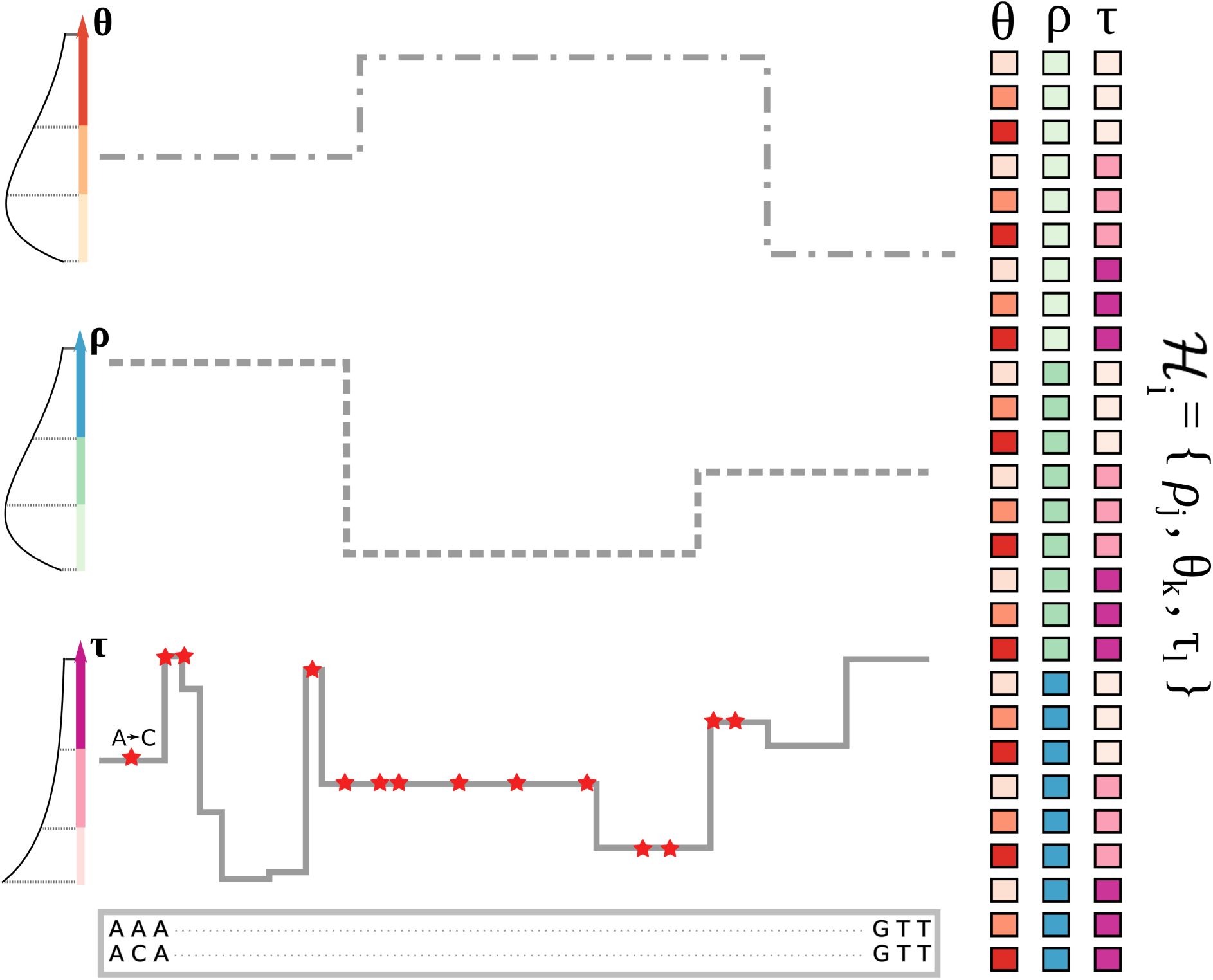
Schematic representation of ρ-θ-iSMC for one pair of genomes. This cartoon model has three time intervals, three recombination rate categories and three mutation rate categories. The genome-wide distribution of diversity depends on the mutation landscape (top) and on the τ landscape (bottom), which is modulated by the recombination landscape (middle). Discretized values of these distributions (left) are combined in triplets as the hidden states of our Hidden Markov Model (right).

The full ρ-θ-iSMC model remains parsimonious, being characterized by a total of 11 parameters, namely, 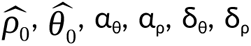 plus five parameters describing constrained cubic splines that embody the demographic curve over time (Barroso et al., 2019). (Such parsimony is afforded by the structure of the Markov-modulated HMM which readily leverages physical linkage among sites in the same chromosome to fit distributions of recombination rates, mutation rates and TMRCA that are shared throughout the genome, even if site-specific realizations of these values may differ.) Running its forward recursion independently on each pair of genomes gives the composite likelihood of the model. After parameter optimization, we seek to reconstruct single-nucleotide landscapes (ρ, θ or τ) for each diploid separately. We first compute the posterior probability of each hidden state for every site *i* in the diploid genomes using regular HMM procedures (Durbin et al., 1998). Afterward, since in ρ-θ-iSMC the hidden states are triplets (**Figure 1**), computing the posterior average of each landscape of interest amounts to first marginalizing the probability distribution of its categories and then using it to weight the corresponding discretized values (Barroso et al., 2019). Let 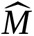 be the inferred discretized Gamma distribution shaping mutation rate variation, and 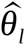 be the product of the estimated genome-wide average mutation rate 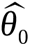 and the value of 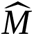 inside category *l*. Similarly, let 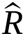 be the inferred discretized Gamma distribution shaping recombination rate variation, and 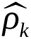 be the product of the estimated genome-wide average recombination rate 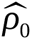 and the value of 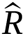 inside category κ. Then the posterior average 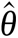 at position *i* is given by:

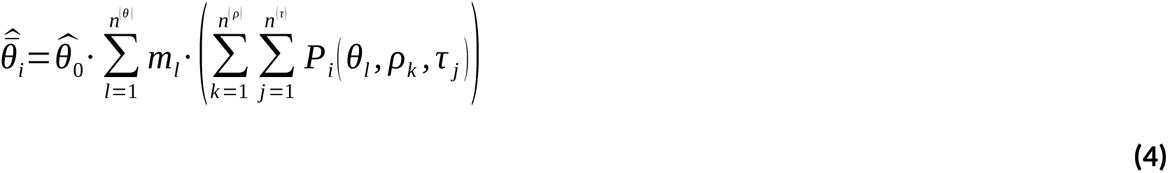

where 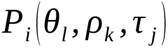 is the probability of the triplet 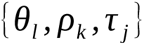 (which denotes a unique hidden state of the model) underlying the *i*-th site of the genome. Likewise, the posterior average 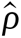 at position *i* is given by:

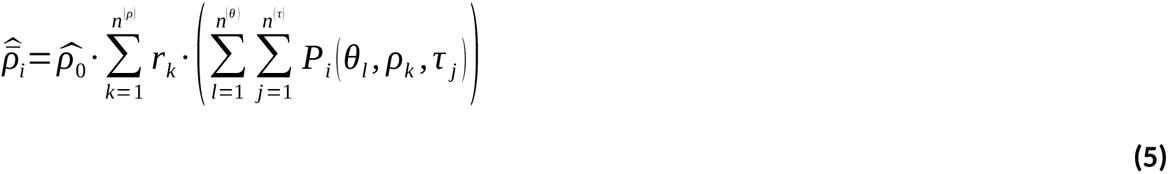

Finally, the posterior average 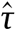 at position *i* is presented in units of _4 *× Ne*_ generations and obtained with:

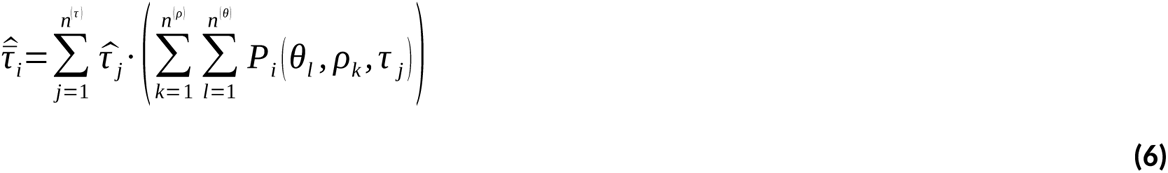

For each diploid, we can then bin the inferred single-nucleotide landscapes into non-overlapping windows of length *L* by averaging our site-specific estimates over all sites within each window. A consensus map of the population is obtained by further averaging over all *n* individual (binned) maps in our sample, i.e.:

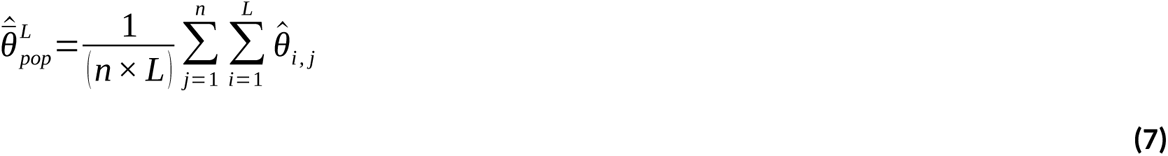

is our estimate of the consensus mutation rate in a single genomic window of length *L*, where *n* is the number of pairs of genomes analyzed by iSMC, and likewise for ρ and τ:

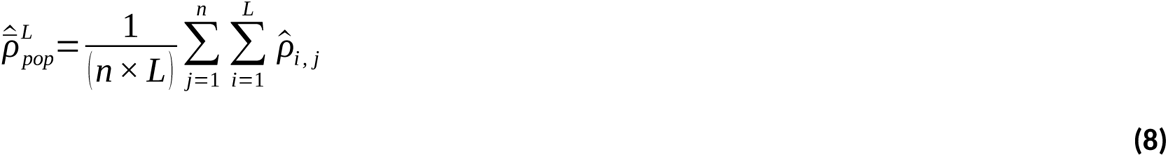

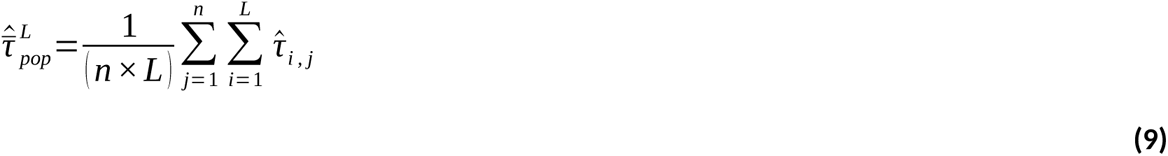

We finally note that the auto-correlation parameters δ_θ_ and δ_ρ_ represent the probabilities of switching mutation and recombination rates between adjacent sites, averaged along the genome. That is, although we include two layers of complexity in comparison to the original SMC models, we assume here that such transition probabilities are themselves spatially homogeneous. In reality, genomic regions may differ in the rate of change between local mutation and recombination rates. Nevertheless, in practice, the reconstruction of mutation and recombination maps with posterior decoding should be somewhat robust to this model mis-specification.

### Simulation study

Using SCRM (Staab et al., 2015), we simulated 10 haploid sequences of length 30 Mb with parameters based on those inferred from ρ-θ-iSMC in *D. melanogaster* (see Results): θ = 0.0112; ρ = 0.036; α_θ_ (continuous Gamma distribution used as mutation rate prior) = 3.0; α_ρ_ (continuous Gamma distribution used as recombination rate prior) = 1.0; δ_θ_ (mutation rate transition probability) = 1e-05; δ_ρ_ (recombination rate transition probability) = 1e-04. Note that such transition probabilities lead to landscapes where blocks of constant mutation and recombination span, on average, 10 kb and 100 kb, respectively, with stochastic variation coming from the geometric distributions used to model them. **Supplemental Figure S1** displays a sketch of the smoothed demographic history used in the coalescent simulations (see Results). **Figures 4** and **5** display the mean R^2^ value of the ANOVA performed on the inferred landscapes from 10 simulated replicates (see Results), but the standard deviation of these estimates are very small, and confidence intervals were, therefore, omitted. Data leading to **Figure 5** was also simulated with SCRM, with parameters described in the Results section.

Next, we used SLiM 3.00 (Haller & Messer, 2018) to simulate the genealogy of a chromosome undergoing purifying selection, using *D. melanogaster*’s chromosome 2L as a template. The simulated region was 23.51 Mb long, and we used Comeron’s recombination map in 100 kb windows (Comeron et al., 2012). We used Ensembl (Cunningham et al., 2022) release 103 gene annotations for *D. melanogaster* and extracted all exons coordinates, merging overlapping exons. Forward simulations were conducted using SLiM, with only deleterious mutations in exons being modeled. The fitness effect of mutations was drawn from a negative gamma distribution with a shape of 1.0 and a mean of –5/10,000. The population size was kept constant and equal to 10,000 and the population evolved for 700,000 generations. To compensate for the low population size, we scaled the mutation and recombination rates by a factor of 10 to result in a scenario closer to the *D. melanogaster* demography. The deleterious mutation rate was set to 1e-7 bp^-1^ along the genome. Ten replicates were generated and saved as tree sequences (Kelleher et al., 2018), which were then further processed by the ‘pyslim’ python module to run a recapitation procedure to ensure that all lineages coalesced into a single root at all genome positions. Ten genomes were then sampled uniformly at random and the underlying tree sequence exported. Finally, ‘msprime’ (Kelleher et al., 2016) was used to add neutral mutations to the tree sequence and save the resulting sequence alignments. A random mutation rate map was generated by sampling relative rates from a Gamma distribution with mean equal to 1.0 and with a shape parameter equal to 2.5, in segments with lengths drawn from a geometric distribution with mean equal to 100 kb. The resulting mutation relative rate map was then scaled by the genome average mutation rate of 1e-7 bp^-1^.

### Analyses of *Drosophila* data

Model fitting and posterior decoding by ρ-θ-iSMC in *D. melanogaster* data were performed using a hidden-states configuration of 30 τ intervals, five ρ categories and five θ categories. We used publicly available data – haplotypes ZI103, ZI117, ZI161, ZI170, ZI179, ZI191, ZI129, ZI138, ZI198 and ZI206 coming from the Zambia population in the Drosophila Population Genomics Project Phase 3 (Lack et al., 2015). Note that the following filters have been previously applied to these data by the original authors: A) heterozygous regions (maintained in the inbred individuals by selection due to recessive lethal alleles); B) three bp around called in-dels; C) long identity-by-descent stretches between genomes from the same location; as well as D) segments showing evidence of recent admixture (from outside Africa back into Africa) were all masked (turned to ‘N’ in the FASTA files). We assigned gaps and masked nucleotides in these FASTA sequences as “missing” data (encoded by the observed state ‘2’ within iSMC, for which all hidden states have emission probability equal to 1.0 (Li & Durbin, 2011)). To optimize computational time, ρ-θ-iSMC was first fitted to chromosome 2L only. Maximum likelihood estimates from this model were then used to perform posterior decoding on all other autosomes. Prior to fitting the linear models, for each scale in which the iSMC-inferred landscapes were binned (50 kb, 200 kb and 1 Mb), we filtered out windows with more than 10% missing data in the resulting maps. Genomic coordinates for coding sequences and their summary statistics (π_N_, and π_S_) were taken from (Moutinho et al., 2019).

### Linear modeling

Linear models implementing our causal model of diversity (**Figure 3**) were built based on genomic maps of 50 kb, 200 kb and 1 Mb resolution. It is worth reiterating that the binning of the singlenucleotide landscapes happens after optimization by the HMM such that it does not influence model complexity (as detailed in the model description, the 11 iSMC parameters are jointly estimated for the entire dataset, i.e., the model is aware of all individual sites in the sequences during optimization). When building linear models from real data, we first fitted GLS models independently to each autosome arm (2L, 2R, 3L, 3R), correcting for both auto-correlation of and heteroscedasticity of the residuals. After using Bonferroni correction for multiple testing, we observed (across the autosome arms and for different window sizes) significant and positive effects of θ and τ on π, whereas the effect of ρ was only significant for chromosome 3L at the 200 kb scale, and the interaction between θ and τ is positive and significant except for arms 2R and 3L at the 1 Mb scale (**Supplemental Tables S6, S7, S8**). Since the trends in coefficients are overall consistent, we pulled the autosome arms and in the Results section we present linear models fitted to the entire genome, for ease of exposition. Because we cannot rely on the GLS to partition the variance explained by each variable using type II ANOVA, we used OLS models to compute R^2^ and restricted the GLS to assess the sign and significance of variables. We standardized all explanatory variables (subtracted the mean then divided by the standard deviation) before fitting the regression models to aid in both computation of variance inflation factors and interpretation of the coefficients.

## Results

### The sequentially Markov coalescent with heterogeneous mutation and recombination

The sequentially Markovian Coalescent (SMC) frames the genealogical process as unfolding spatially along the genome (Marjoram & Wall, 2006; McVean & Cardin, 2005; Wiuf & Hein, 1999). Its first implementation as an inference tool derives the transition probabilities of genealogies between adjacent sites as a function of the historical variation in *N*_e_ (i.e., demographic history) and the genome-wide average scaled recombination rate *_ρ_*_=4 *× Ne ×r*_ (Li & Durbin, 2011). Model fitting is achieved by casting the SMC as a hidden Markov model (HMM) (Dutheil, 2017) and letting the emission probabilities be functions of the underlying Time to the Most Recent Common Ancestor (TMRCA, τ) and the scaled mutation rate *_θ_*_=4 *× Ne ×μ*_ (see Methods). The SMC has proven to be quite flexible and serves as the theoretical basis for several models of demographic inference (see Spence et al. (2018) for a review, and Sellinger et al. (2020) for another compelling, more recent development). We have previously extended this process to account for the variation of ρ along the genome, thereby allowing for a heterogeneous frequency of transitions between local genealogies in different parts of the genome (Barroso et al., 2019). In this more general process called iSMC, recombination rate heterogeneity is captured by an autocorrelation parameter, δ_ρ_, where the localized values of ρ are taken from a discrete distribution and the transition between recombination rates along the genome follows a first-order Markov process.

In the general case, the iSMC process is a Markov-modulated Markov process that can be cast as an HMM where the hidden states are *n*-tuples storing all combinations of genealogies and discretized values of each parameter that is allowed to vary along the genome (Dutheil, 2021). If one such parameter contributes to either the transition or emission probabilities of the HMM, then the hyper-parameters that shape its prior distribution can be optimized, e.g. by maximum likelihood (see Methods). In the iSMC with heterogeneous recombination (ρ-iSMC) the hidden states are pairs of genealogies and recombination rates (Barroso et al., 2019). Here, we extend this model by allowing the mutation rate to also vary along the genome (**Figure 1**), following an independent Markov process, i.e., letting the hidden states of the HMM be {θ-category, ρ-category, genealogy} triplets. The signal that spatial variation in ρ and θ leaves on the distribution of SNPs is discernible because their contributions to the likelihood are orthogonal: the recombination and mutation rates affect the transition and emission probabilities of the forward HMM algorithm, respectively. Parameter optimization and subsequent posterior decoding is performed as in Barroso et al. (2019). Under strict neutrality (which results in *N*_e_ being homogeneous along the genome (Charlesworth, 2009)), the inferred θ landscape reflects the landscape of *de novo* mutations (μ). iSMC can, therefore, be used to infer genome-wide variation in mutation rates with single-nucleotide resolution and statistical noise is reduced by averaging the posterior estimates of θ within larger genomic windows (see Methods).

In order to increase power, we further extend iSMC to accommodate multiple haploid genomes. In this augmented model, input genomes are combined in pairs such that the underlying genealogies have a trivial topology reduced to their τ (Figure 1). Although under Kingman’s Coalescent (Kingman, 1982) the genealogies of multiple pairs of genomes are not independent, we approximate and compute the composite log-likelihood of the entire dataset by summing over such “diploid” log-likelihoods, similarly to MSMC2 (Malaspinas et al., 2016). Furthermore, iSMC enforces all diploids to share their prior distributions of τ, ρ and θ so that multiple sequences provide aggregate information to our parameter inference during model fitting; it does not, however, explicitly enforce that they have identical genomic landscapes upon posterior decoding. Rather, iSMC uses posterior probabilities to reconstruct recombination and mutation maps separately for each diploid.

Especially at the single-nucleotide level, accuracy of the inferred posterior landscapes is limited by the large stochasticity of the coalescent (Hein et al., 2004). The combination of genealogical and mutational variance leads to differences among the posterior landscapes of θ and ρ inferred from each diploid because it creates departures from the expected number of SNPs along pairs of genomes (hence variation in the amount of information diploids bear, in different regions of the genome, about ancestral processes such as mutation and recombination). To reduce noise from the individual diploid estimates and obtain consensus landscapes of the whole sample, iSMC averages the posterior estimates of θ and ρ over all diploids, for each site in the genome (see Methods). On the other hand, differences in the τ landscapes among diploids primarily reflect the stochastic nature of the ancestral recombination graph along the genome, which has intrinsic value itself. We therefore average these diploid τ landscapes not to reduce estimation noise but to obtain a measure of drift in neutral simulations. Note, however, that the average τ of the sample within a genomic window also contains information about natural selection (Palamara et al., 2018) – a property we exploit in the analyses of *Drosophila* data.

### Mutation rate variation impacts nucleotide diversity more than linked selection in *Drosophila*

We sought to quantify the determinants of genome-wide diversity in *D. melanogaster* using 10 haploid genome sequences from the Zambia population. To infer the genomic landscapes, we employed a ρ-θ-iSMC model with five mutation rate classes, five recombination rate classes and 30 coalescence time intervals, leading to 750 hidden states. We note that the number of classes and time intervals do not affect the number of estimated parameters, in particular because our implementation of the demographic model uses splines in place of the emblematic “skyline” model (Li & Durbin, 2011; Schiffels & Durbin, 2014) (see Methods). In general, finer discretization of these three distributions leads to more precise inference until a plateau is reached, as well as impacts the minimum and maximum values that the posterior estimates can take. However, the memory use and the likelihood computation time scale linearly and quadratically with the total number of hidden states, respectively. We selected 30 classes for the TMRCA and five classes for each rate distribution because this configuration provided a good tradeoff between computational resources and accuracy during our testing phase. The total run-time for fitting the model with 750 hidden states to chromosome 2L of *D. melanogaster* was about 1 month on a highperformance cluster. Therefore, we proceeded in two steps: we first estimated model parameters on a subset of the data (chromosome arm 2L), and then used the fitted model to infer the landscape of mutation, recombination and TMRCA for all autosomes (see Methods). The justification for this approach is that the HMM posterior decoding is able to reconstruct chromosome-specific landscapes, even from identical prior distributions. At the same time, we have no *a priori* reason to believe that the shape of these distributions will differ substantially among autosomes. The similarity among the results obtained with each chromosome in the downstream analyses (see “Linear Modeling” sub-section within Methods) supports such intuition (**Supplemental Tables S6, S7** and **S8**).

The iSMC parameters estimated from *D. melanogaster* suggest an exponential-like distribution of recombination rates (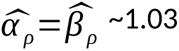 for their Gamma distribution) whereas the inferred distribution of mutation rates is more tightly centered around the mean (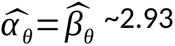 for their Gamma distribution). iSMC also inferred that the change in recombination rate across the genome was more frequent (autocorrelation parameter 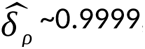, corresponding to a change of recombination rate on average every 10kb) than the change in mutation rate (auto-correlation parameter 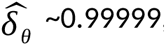, corresponding to a change of mutation rate on average every 100 kb). This suggests that our model mostly captures largescale rather than fine-scale variation in the mutation rate. Our inferred genome-wide average 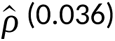 is in line with previous estimates (Chan et al., 2012), and the coalescence rates (which, in the context of this article, comprise a collection of nuisance parameters used to refine our estimates of τ, ρ and θ along the genome) suggest a ∼4-fold bottleneck followed by recovery (**Supplemental Figure S1**). As an empirical validation of this new iSMC method, Spearman’s rank correlations (hereafter referred to as Spearman’s rho) between our inferred recombination map of chromosome 2L and Comeron’s map based on experimental crosses (Comeron et al., 2012) are 0.594 at the 50 kb scale, 0.693 at the 200 kb scale and 0.865 at the 1 Mb scale (all p-values < 1e-5), higher than the correlations reported with previously published population genetic methods applied to *D. melanogaster* (Adrion et al., 2019; Barroso et al., 2019; Chan et al., 2012).

We used the parameters estimated from *D. melanogaster* to simulate 10 replicate datasets under a purely neutral scenario (see Methods). The aims of these simulations are two-fold: (1) to benchmark iSMC’s accuracy in reconstructing the mutation landscape; and (2) to understand how ρ, θ and τ interact to influence diversity levels under neutrality, thereby providing a measure of contrast for the analyses of real data (where natural selection is present). Throughout this article, we analyze the determinants of nucleotide diversity at different scales by binning the landscapes of mutation, recombination and TMRCA into non-overlapping windows of 50 kb, 200 kb and 1 Mb. We first report strong correlations between inferred and simulated maps, ranging from 0.975 to 0.989 (Spearman’s rho, **Figure 2A**, **Supplemental Table S1**), showcasing that our model is highly accurate under strict neutrality and when mutation rate varies along the genome in Markovian fashion.

**Figure 2.**
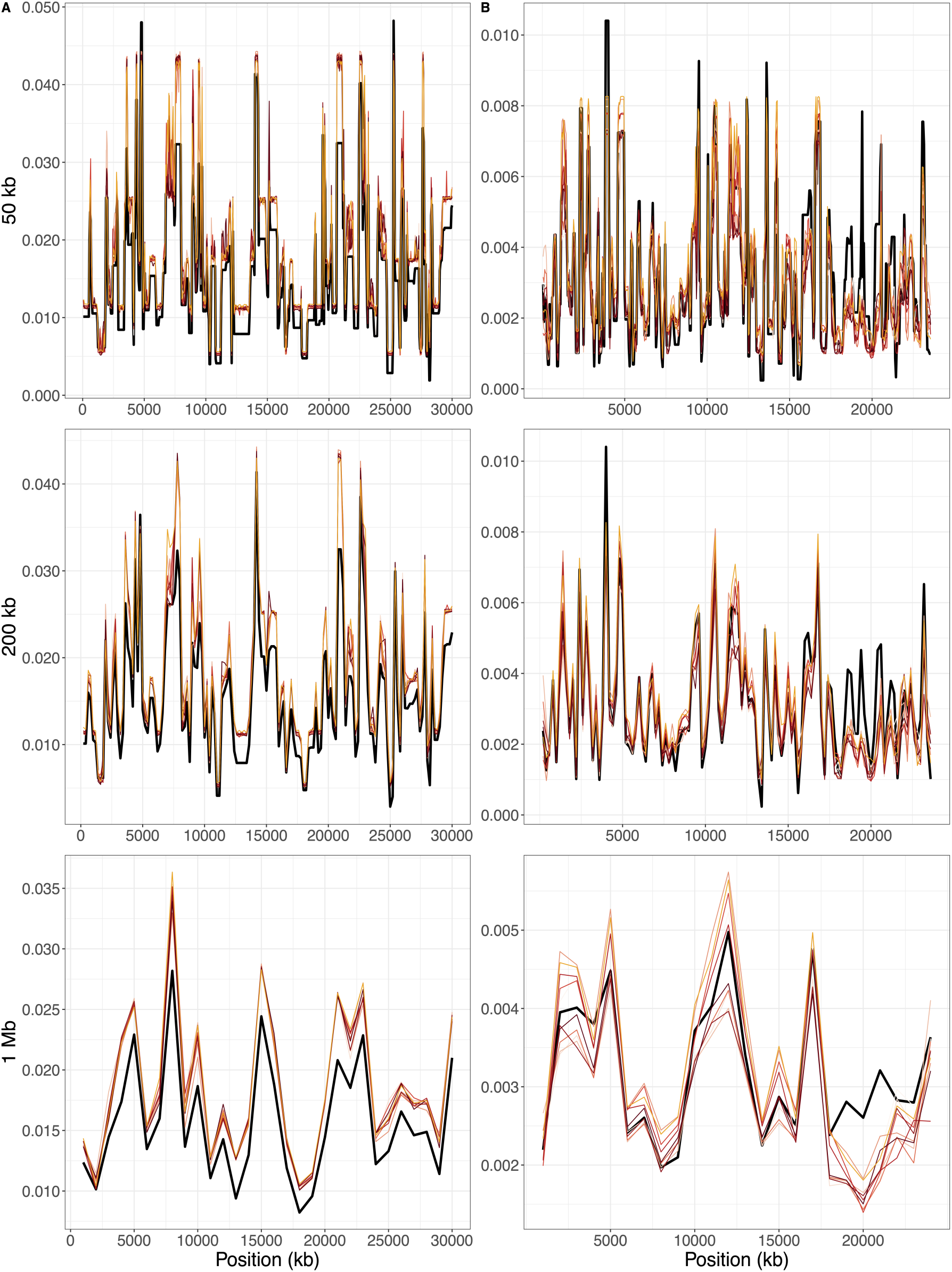
iSMC recovers the mutation landscape in simulations. A) Coalescent simulations under neutrality. **B)** Simulations with background selection. In both cases, the simulated mutation landscape is shown by the thick black line whereas inferred landscapes are shown, for each replicate, by thin lines in shades of red. From top to bottom: 50 kb scale, 200 kb scale, 1 Mb scale.

**Table 1.** Estimates from linear regression models fitted to the distribution of nucleotide diversity along Drosophila melanogaster genomes. Vertical panels show results according to genomic window size whereas horizontal panels show results according to the structure of the linear model. OLS = Ordinary Least Squares; GLS = Generalized Least Squares; VIF = Variance Inflation Factor.

| Model | Type | Variable | 50 kb |  |  | 200 kb |  |  | 1 Mb |  |  |
| --- | --- | --- | --- | --- | --- | --- | --- | --- | --- | --- | --- |
|  |  |  | Coefficient | p-value | VIF | Coefficient | p-value | VIF | Coefficient | p-value | VIF |
| 1 | OLS | $\theta$ | 0.0027 | <2.2e-16 | 1.3 | 0.0026 | <2.2e-16 | 1.7 | 0.0025 | <2.2e-16 | 2.4 |
| | | $\tau$ | 0.0010 | <2.2e-16 | 2.5 | 0.0008 | <2.2e-16 | 6.0 | 0.0006 | <2.2e-16 | 13.0 |
| | | $\rho$ | 0.00004 | 0.0788 | 1.5 | 0.00001 | 0.2110 | 1.7 | 0.0000003 | 0.0350 | 1.8 |
| | | $\theta:\tau$ | 0.0003 | <2.2e-16 | 1.6 | 0.0002 | <2.2e-16 | 3.7 | 0.0001 | <1e-4 | 7.7 |
| 2 | GLS | $\theta$ | 0.0027 | <1e-4 | 1.2 | 0.0026 | <1e-4 | 1.4 | 0.0024 | <1e-4 | 1.6 |
| | | $\tau$ | 0.0010 | <1e-4 | 1.9 | 0.0008 | <1e-4 | 3.7 | 0.0007 | <1e-4 | 1.8 |
| | | $\rho$ | 0.000004 | 0.3897 | 1.4 | 0.000005 | 0.5720 | 1.5 | 0.000003 | 0.7980 | 1.8 |
| | | $\theta:\tau$ | 0.0003 | <1e-4 | 1.3 | 0.0002 | <1e-4 | 2.5 | 0.0001 | 0.0003 | 1.3 |
| 3 | GLS | $\theta$ | 0.0030 | <1e-4 | 1.0 | 0.0029 | <1e-4 | 1.0 | 0.0027 | <1e-4 | 1.1 |
| | | $\rho$ | 0.00040 | <1e-4 | 1.0 | 0.0003 | <1e-4 | 1.0 | 0.0002 | <1e-4 | 1.1 |

We then used the raw genomic landscapes from these simulated (neutral) datasets to investigate how evolutionary mechanisms shape the distribution of nucleotide diversity along the genome, measured as π, the average per-site heterozygosity of the sample. The structure of our hypothesized causal model of diversity (solid lines in **Figure 3**) is rid of “backdoor paths” that would otherwise create spurious associations between recombination, mutation, TMRCA and nucleotide diversity (Pearl & Mackenzie, 2018). We could thus cast our causal model as an ordinary least squares regression (OLS) that seeks to explain π as a linear combination of the standardized variables ρ, θ and τ and statistical associations between our explanatory variables and the outcome variable π then represent causal relationships that merit scientific explanation. The justification for a linear model of π is that for sufficiently small genomewide average diversity θ_0_ (a requirement which is met in *D. melanogaster*, as 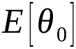 ~ 1e-2) the per-site heterozigosity 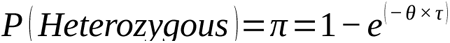 can be well approximated by 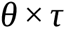 the first term in the Taylor series expansion of 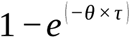. Since simulations grant direct access to the true genomic landscapes, then by definition the ensuing OLS models are free of estimation noise in the explanatory variables and serve as a ground truth assessment of how neutral evolutionary mechanisms influence nucleotide diversity. Because of the interplay between genealogical and mutational variance, we tested the improvement that including an interaction term between θ and τ brought to the fit of the linear models. In all replicates, we found that model selection using Akaike’s information criterion favors a regression with an interaction term between the two variables that directly influence nucleotide diversity, 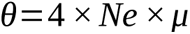 over the simpler model 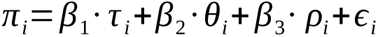.

**Figure 3.**
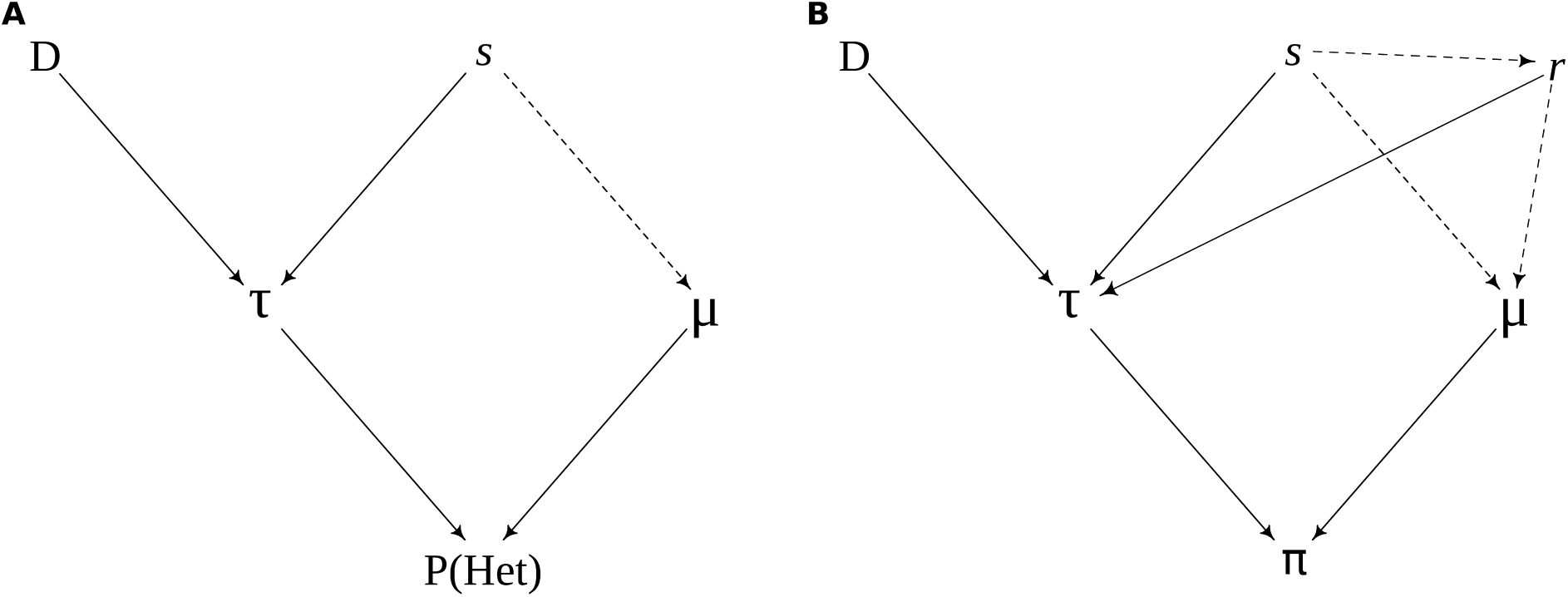
Directed acyclic graphs depicting our abstract causal model for the determinants of genome-wide diversity. A) for a single, hypothetical nucleotide that is independent of any neighbors, its probability of being heterozygous is solely influenced by the local mutation rate (μ) and TMRCA (τ), which in turn is affected by drift (D) and selection (s). **B)** when contiguous sites are grouped into genomic windows, their correlated histories imply that the local recombination rate (r) plays a role in modulating both D and the breadth of linked selection via τ, which together with local μ influences π. Relationships that may be relevant in other model systems are shown by dashed lines (where selection affects μ and r through modifier genes and where recombination is mutagenic). Note that P(het) in **A** has exactly the same form as the emission probability of the HMM model, 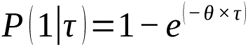.

Fitting the regression model at the 50 kb, 200 kb and 1 Mb scales shows significant and positive effects of θ and τ, but not of ρ, on π (**Supplemental Table S2**, upper panel). This is expected since both deeper ancestry and higher mutation rate lead to increased nucleotide diversity and the influence of recombination rate on π is mediated by τ, thus disappearing due to its inclusion in the linear model. There is also a significant and positive effect of the interaction between θ and τ, highlighting the interplay between genealogical and mutational variance, where the effect of the mutation rate on diversity can only be fully manifested if ancestry is deep enough (reciprocally, ancestry can only be seen clearly if the local mutation rate is high enough). Moreover, the standardization that we employed on the explanatory variables prior to fitting the linear models (see Methods) allows us to evaluate their relative importance to the π distribution straight from the estimated coefficients. We observe that the linear coefficient of θ is ∼6 times larger than the linear coefficient of τ at the 50 kb scale, ∼11 times larger at the 200 kb scale and ∼16 times larger at the 1 Mb scale (**Supplemental Table S2**, upper panel). Besides the linear coefficients, we further quantified the relative influence of mutation, drift and recombination to local diversity levels by partitioning the R^2^ contributed by each explanatory variable with type II ANOVA. Consistently with the previous results, our estimates show that the θ landscape explains most of the variance in π in our simulations and that its contribution increases with the genomic scale (96.3% at 50 kb, 98.6% at 200 kb and 99.3% at 1 Mb **Figure 4A**). On the other hand, the contribution of the τ landscape decreases with the genomic scale (2.7% at 50 kb, 1% at 200 kb and 0.54% at 1 Mb). We propose that these trends stem from the minuscule scale of variation in τ (changing on average every 48.42 bp due to recombination events in our coalescent simulations, median = 19 bp), which smooth out more rapidly than does mutation variation when averaged within larger windows. Conversely, the broader scale of heterogeneity in θ (changing every 100 kb on average) makes it comparatively more relevant at larger window sizes. Strikingly, the total variance explained by the model is >99% at all scales, suggesting that these three landscapes are sufficient to describe the genome-wide distribution of diversity, as illustrated by our causal model (**Figure 3**). To test whether we could recover such trends with the landscapes inferred by our HMM, we fitted the OLS models to the same genomic landscapes of nucleotide diversity except using the maps inferred by iSMC as explanatory variables (i.e., 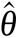, 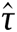 and 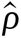 instead of the true, simulated ones: θ, τ and ρ). The sign and significance of the estimated OLS coefficients remained unchanged (**Supplemental Table S2**, middle panel), as do the ranking of their effect sizes, but in some replicates the residuals of the model were found to be correlated and/or with heterogeneous variance. As this violation of the OLS assumption could bias the estimates of the p-values of the linear coefficients, we also fitted Generalized Least Squares (GLS) models accounting for both deviations, which reassuringly produced coherent results (**Supplemental Table S2**, lower panel). Although co-linearity between 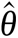 and 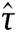 arises due to confounding in their estimation by iSMC, the variance inflation factors are always < 5, indicating that the coefficients are robust to this effect (Ferré, 2009). The trends in the linear coefficients obtained with iSMC-inferred landscapes are the same as those obtained with simulated (noise-free) landscapes, except that the effect of 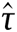 is estimated to be larger than that of τ. Similarly, type II ANOVA using the inferred landscapes shows that the contribution of 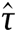 is slightly higher than when using the true landscapes (5.1%, 2.9% and 1.4%, increasing window size) whereas the contribution of 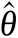 is slightly lower (92.5%, 95.4% and 97.5%, increasing window size), but the variance explained by each variable closely agrees between the two cases (middle and right panels in **Figure 4A**). Therefore, we conclude that the joint-inference approach of iSMC can infer the genomic landscapes of τ, ρ and θ and that the linear regression representation of our causal model (**Figure 3**) is able to quantify their effect on the distribution of nucleotide diversity, π.

**Figure 4.**
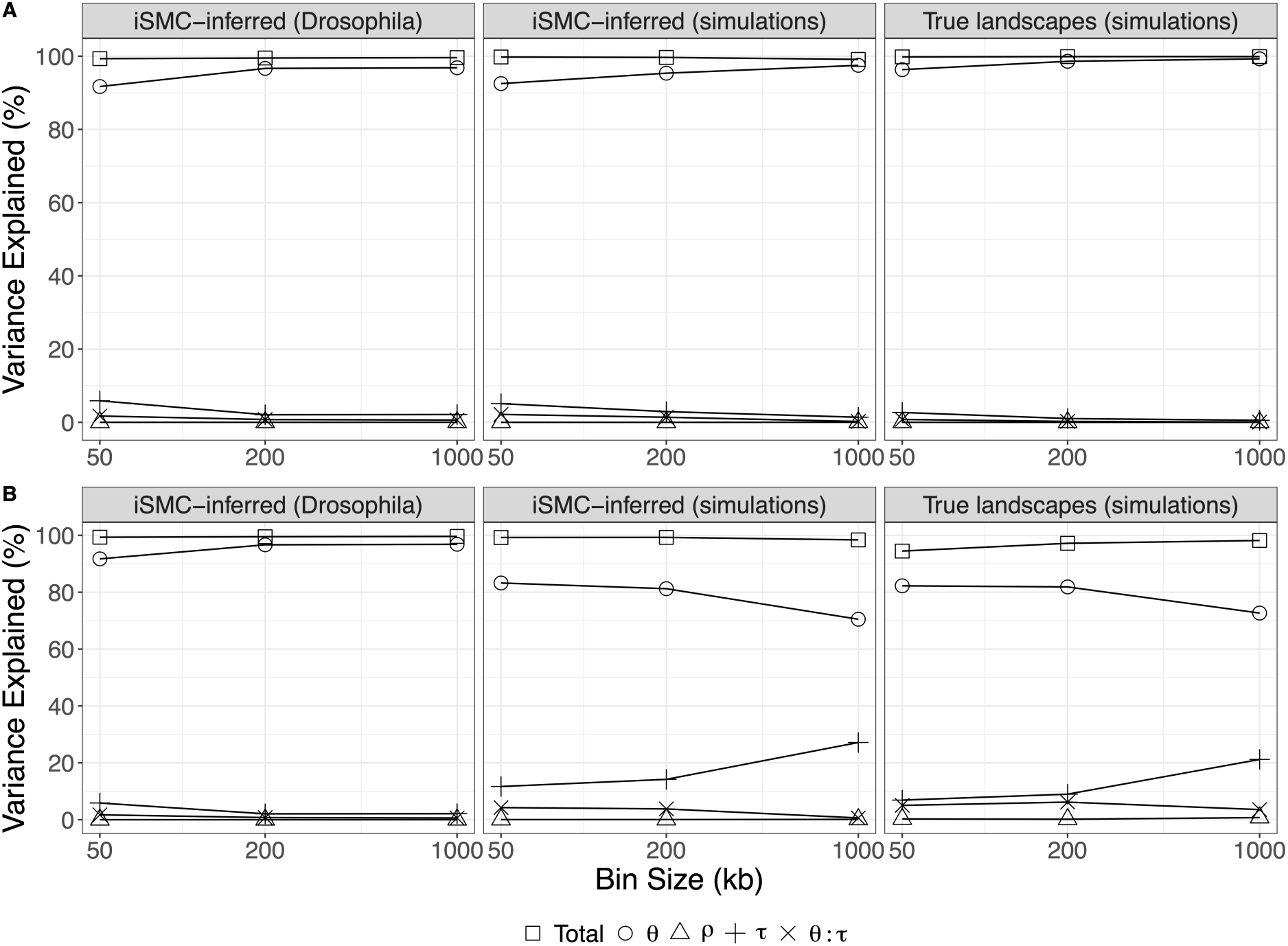
Variance in the distribution of diversity explained by each genomic landscape. Partitioning of variance according to window size (x-axis, shown in log_10_ scale), using either simulated data (true landscapes: right panels; inferred landscapes: middle panels) or real *Drosophila* data (left panels). **A)** comparison between real *Drosophila* data and results from neutral simulations. **B)** comparison between real *Drosophila* data and results from simulations under background selection. In each panel, shapes represent explanatory variables in the linear model: θ (circles), ρ (triangles), τ (plus sign), θ:τ interaction (crosses) and the total variance explained by the model (squares) is the sum of the individual R^2^. Each point represents the average R^2^ over 10 replicates. Variation among replicates resulted in confidence intervals too small to be plotted.

We finally employed the landscapes obtained with ρ-θ-iSMC to quantify the determinants of genomewide diversity in *D. melanogaster*. In the following analyses, our interpretations of the OLS models assume that sequencing errors are unbiased with respect to the explanatory variables and that the population is broadly panmictic (or that geographic structure is implicitly accounted for by the TMRCA, e.g. (Beichman et al., 2018)). We also follow previous work suggesting that recombination is not mutagenic in this system (Begun et al., 2007; Castellano et al., 2016; McGaugh et al., 2012), thus we ignore this potential relationship. We used our inferred *D. melanogaster* maps to fit an OLS regression of the form 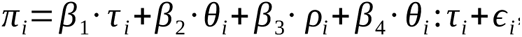 As in our simulations, the regression model shows positive effects of both 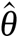 and 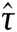 but not of 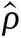 on π across all scales (**Table 1**). Likewise, a GLS model correcting for the identified auto-correlation of and heteroscedasticity of the residuals yields the same trends, and its variance inflation factors are < 5, indicating that the estimated coefficients are robust to co-linearity (Ferré, 2009). Showcasing its dominant impact on π in the fruit fly, the linear coefficient of 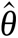 is between three and four times larger than that of 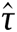 a trend that is akin to that obtained with inferred maps in the coalescent simulations. Moreover, partitioning of variance shows a small contribution of 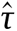 that decreases with increasing genomic scale (5.9% at 50 kb, 2.1% at 200 kb and 2.1% at 1 Mb) whereas the opposite applies to 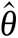 (91.7% at 50 kb, 96.7% at 200 kb and 96.8% at 1 Mb, left panel in **Figure 4A**). Our linear model explains >99% of the variation in π along *D. melanogaster* autosomes, and the effects of our inferred landscapes on diversity are remarkably close to those from our neutral simulations (**Figure 4A**), suggesting that iSMC is robust to the occurrence of selection in this system. Unlike neutral simulations; however, the simple correlation test between 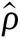 and π ends up positive and significant in *D. melanogaster* data, at least at smaller scales (Spearman’s rho = 0.20, p-value = 2e-13 at the 50 kb scale; Spearman’s rho = 0.15, p-value = 0.0025 at the 200 kb scale; Spearman’s rho = 0.20, p-value = 0.07 at the 1 Mb scale), recapitulating the classic result of Begun & Aquadro (1992) and indicating the presence of linked selection. We also found a positive correlation between 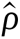 and 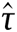 (Spearman’s rho = 0.48, p-value < 2.2e-16 at 50 kb; Spearman’s rho = 0.45, p-value < 2.2e-16 at 200 kb; Spearman’s rho = 0.48, p-value < 2.2e-16 at 1 Mb), once again contrasting the results under neutrality and suggesting that the effect of linked selection is indeed captured by the distribution of genealogies and modulated by the recombination rate (Cutter & Payseur, 2013). Although τ is primarily influenced by demography in SMC-based models (by means of a Coalescent prior taming the transition probabilities of the HMM (Li & Durbin, 2011; Schiffels & Durbin, 2014), it has also been demonstrated to carry the signature of selection due to local changes in coalescence rates that have been interpreted as spatial variation in *N*_e_ (Palamara et al., 2018; Zeng & Charlesworth, 2011). Shortly, Palamara’s ASMC method reconstructs the TMRCA landscape of several pairs of genomes and interprets recurrent (shallow) outliers in the 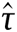 distribution as the outcome of linked selection (i.e., regions where pair of genomes consistently coalesce faster than expected under neutrality). We tested the sensitivity of our regression framework to this effect by a fitting linear model without 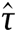 as an explanatory variable, 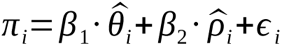 hypothesizing that in the absence of its mediator the recombination rate would show a significant and positive effect on diversity. Indeed, this is what we found at all genomic scales (**Table 1**, Model 3), corroborating our interpretation of the causal relationships in the presence of selection (**Figure 3B**), from which the direct correlation between 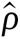 and π, often reported in the literature, is a special case. In summary, our results show that recombination shapes diversity via the τ distribution and linked selection, but that in *D. melanogaster*, the impact of genetic hitchhiking on the diversity landscape is smaller than that of mutation rate variation.

To investigate the signature of selection, we analyzed the relationship between the local mutation rate and the levels of synonymous (π_S_) and non-synonymous (π_N_) diversity across *D. melanogaster* genes (see Methods). We computed these summary statistics across exons and matched their coordinates with our finest (50 kb-scale) genomic landscapes to increase resolution (i.e., to maximize variation in mutation and recombination rates among genes). We observed a stronger relationship between 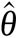 and π_S_ (Spearman’s rho = 0.68, 95% CI after 10,000 bootstrap replicates = [0.64, 0.72], partial correlation accounting for 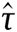) than between 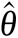 and π_N_ (Spearman’s rho = 0.27, 95% CI after 10,000 bootstrap replicates = [0.22, 0.32], partial correlation accounting for 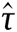) indicating that selection partially purges the excess of non-synonymous deleterious variants in genes with elevated mutation rate, whereas synonymous variants segregate more freely either because they are not directly affected by selection (but are still linked to selected sites) or because selection on codon usage (Lawrie et al., 2013; Machado et al., 2020) is not as strong as selection on protein function. Since synonymous variants are interdigitated with non-synonymous variants, the contrast between these correlation tests cannot be explained by a bias in iSMC’s estimation of θ in functionally constrained regions of the genome. Furthermore, a correlation test between 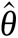 and the proportion of exonic sites in the same 50 kb windows (Spearman’s rho = –0.037, p-value = 0.19, partial correlation accounting for 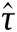) fails to reveal such putative bias (see Discussion for a flip side view on this test). Conversely, we observed a negative and significant correlation between 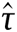 and the proportion of exonic sites (Spearman’s rho = –0.158, p-value = 2e-12, partial correlation accounting for 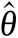), as expected since background selection should reduce the TMRCA more abruptly in densely constrained regions (Charlesworth, 2013; Palamara et al., 2018). We also fitted linear models considering only 50 kb windows with more than 20,000 coding sites. Once again, there were significant and positive effects of both 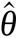 and 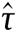 but not of 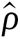, on π. Moreover, the mutation landscape remains the most important factor, explaining 93.2% of the distribution of diversity in gene-rich regions.

### Mutation rate variation shapes genome-wide diversity in neutral scenarios

Our analyses of *D. melanogaster* data and *D. melanogaster*-inspired simulations suggest that the mutation landscape is the main factor influencing levels of diversity along the genome. But are there scenarios where τ has a more pronounced effect on π? We addressed this question by exploring the parameter space of our neutral simulations. For fixed values of the long-term average population size (*N*_e_ = 100,000), the average mutation rate per site per generation (μ = 2e-09), the Gamma distribution of scaling factors of θ (α_θ_ = β_θ_ = 2.5) and the Gamma distribution of scaling factors of ρ (α_ρ_ = β_ρ_ = 1.0), we varied the demographic history (flat *N*_e_; 10-fold bottleneck happening 0.5 coalescent time units ago), the average recombination rate per site per generation (*r* = 1e-08; 1e-09) and the fluctuations of the mutation landscape, where the realized lengths of genomic blocks of constant μ were drawn from geometric distributions with averages equal to 50 kb, 500 kb, or instead taken as a perfectly flat mutation landscape. We reasoned that the extent of the variation in τ along the genome compared to that of μ (equivalently, θ) should modulate their relative influence on π. We fitted OLS models to explain π using the true, simulated landscapes as explanatory variables, and computed their average R^2^ over all replicates for each evolutionary scenario (**Figure 5**). The OLS models included an interaction term between θ and τ but its individual R^2^ was excluded from the plots because it is overall low (∼1%) and of no direct interest. We observed clear trends emerging from these simulated data. First, for a given demographic history and pattern of variation in the mutation rate, increasing *r* reduces the influence of τ on π. This happens because with high recombination rates the genealogies change more often along the genome, thus displaying more homogeneous maps when averaged within windows (50 kb, 200 kb, 1 Mb). Second, for a given *r* and pattern of variation in the mutation rate, τ has a larger impact on π in the bottleneck scenario compared to the scenario of constant population size. This happens because when *N*_e_ varies in time, the distribution of coalescence times becomes multi-modal (Hein et al., 2004) and therefore more heterogeneous along the genome. Third, for a given demographic history and fixed value of *r*, frequent changes in μ along the genome (on average every 50 kb) reduce its impact on π relative to rare changes in μ (on average every 500 kb). This happens because frequent changes in μ lead to more homogeneity along the genome, when averaged within the window sizes used in our analyses.

**Figure 5.**
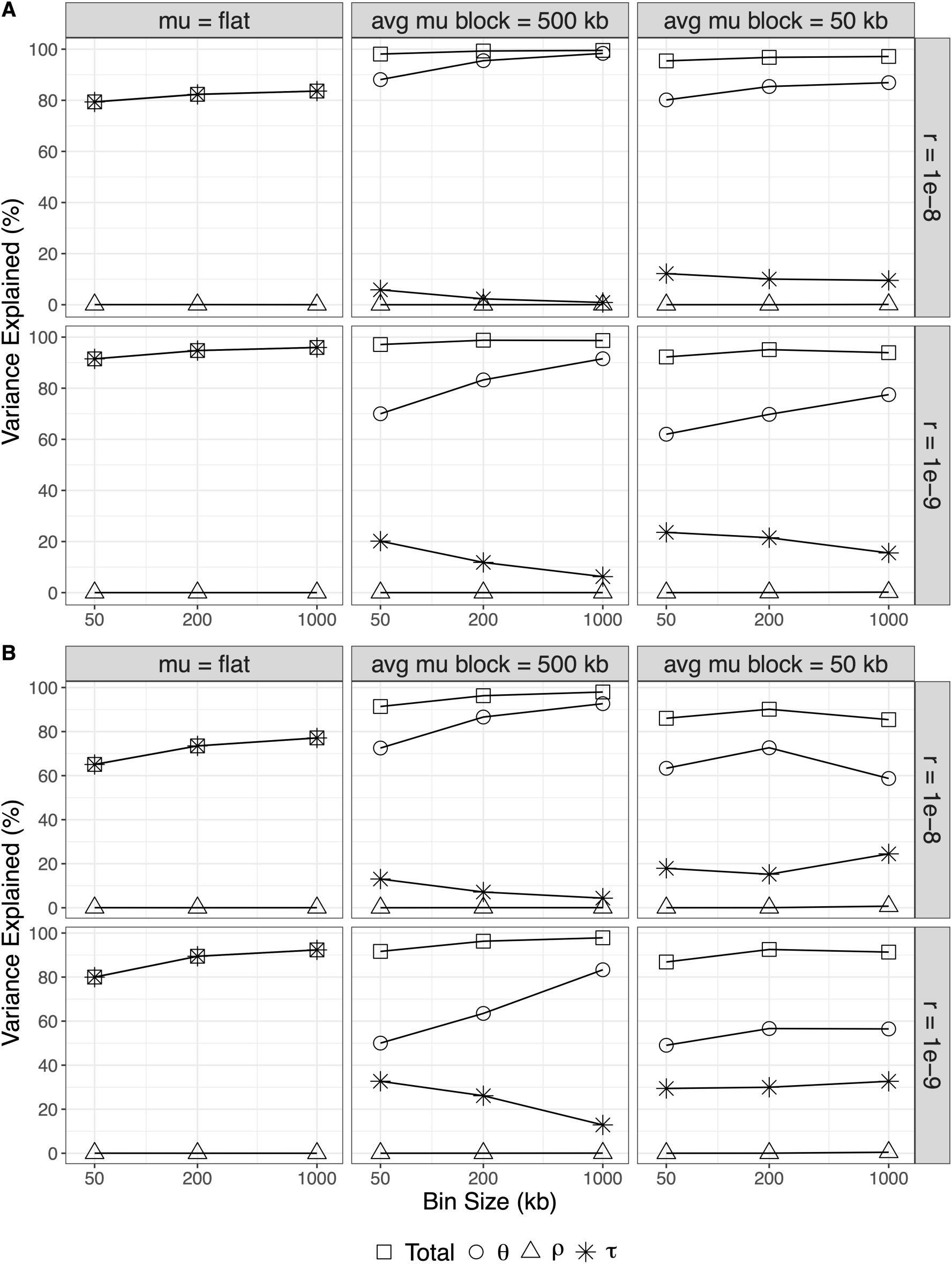
Variance in the distribution of diversity explained by each genomic landscape (neutral simulation study). Partitioning of variance according to window size (x-axis, shown in log_10_ scale). **A)** Constant population size. **B)** Population bottleneck. Results are displayed according to parameters (rows = recombination rate, columns = scale of mutation rate variation). In each panel, point shapes represent explanatory variables in the linear model: θ (circles), ρ (triangles), τ (asterisks) and the total variance explained by the model (squares) is the sum of the individual R^2^. Each point represents the average R^2^ over 10 replicates, and variation among replicates resulted in confidence intervals too small to be plotted. All linear models were built using simulated (true) landscapes.

Finally, if the mutation landscape is flat, then, as expected, the variance explained by our linear model is entirely attributed to τ. Note that although in these neutral simulations τ varies along the genome as a result of genetic drift alone, it still has a non-negligible effect on the distribution of diversity in most scenarios (i.e., binning into large genomic windows does not flatten the TMRCA landscapes completely). This is in agreement with an observation that heterogeneous recombination rates lead to outliers in genome-wide F_ST_ scans, even under neutrality (Booker et al., 2020), which in turn happens because the recombination landscape enlarges the variance of the τ distribution by making the frequency of genealogy transitions a function of the local ρ (confirming the causal effect ρ → τ depicted in **Figure 3B**). From a practical standpoint, it means that drift should not be neglected as an explanation for the distribution of π, especially at narrow window sizes (≤ 10 kb). This is relevant because it is also at narrow window sizes that the effect of selection on diversity levels along the genome can be more easily confounded by the effect of drift, with extreme examples happening during population range expansions and especially in regions of low recombination (Schlichta et al., 2022).

More generally, our simulation study of neutral scenarios shows that the relative impacts of evolutionary mechanisms on π depend primarily on (1) the joint patterns of variation of ρ, τ and θ along the genome; and (2) the window size used in the analysis, because of averaging effects when building the genomic landscapes. In light of these results, the genome of *D. melanogaster* – with its high effective recombination rate, broad (as detectable by iSMC) pattern of variation in the mutation rate and high density of functional sites – seems to be particularly susceptible to the effect of the mutation landscape on its large-scale distribution of diversity. Yet, since the mutation landscape stood out as the most relevant factor in all of the explored (neutral) scenarios where it was allowed to vary (**Figure 5**), we predict that it is likely to shape genome-wide diversity patterns in other species as well.

### iSMC can disentangle mutation rate variation from linked selection

Finally, we simulated 10 replicate datasets under a background selection model with genomic features partially mimicking those of *D. melanogaster* chromosome 2L (see Methods). Briefly, we included in these simulations the positions of exons from the Ensembl database (Cunningham et al., 2022) (whose non-synonymous mutations had selection coefficients drawn from a negative Gamma distribution), the (Comeron et al., 2012) recombination map of the fruit fly, and spatial variation in mutation rates using parameters estimated in our previous analyses. These datasets are not meant to precisely reproduce patterns of nucleotide diversity in real data – there are far too many biological processes not captured by the simulations –, but instead to assess iSMC’s ability to disentangle the θ and τ landscapes when the latter is heavily distorted by linked selection. As such, we used a distribution of selection coefficients with a shape parameter equal to 1.0 (which contrasts with the range from ∼0.3 to ∼0.4 reported in the literature, e.g. Castellano et al. (2018)) in order to artificially exacerbate the effect of linked selection (see below). As before, we fitted a ρ-θ-iSMC model with five mutation rate classes, five recombination rate classes and 30 coalescence time intervals, and afterwards binned the simulated and inferred maps into windows of 50 kb, 200 kb and 1 Mb. We then assessed the accuracy of our framework in two ways: first, by computing Spearman’s rho between simulated and iSMC-inferred landscapes; second, by contrasting the variance explained by each variable in the OLS regression (fitted with inferred versus true maps).

We report strong and positive correlations between simulated and inferred landscapes under such complex a scenario (**Figure 6**). As expected, the inherently fine-scale variation in the distribution of genealogies is the hardest to reconstruct: the Spearman’s rho between the true TMRCA landscapes and the inferred ones ranges from 0.385 to 0.465 at 50 kb and increases substantially with window size (up to 0.787 at 1 Mb, **Supplemental Table S3**). In comparison, the correlation between the true and inferred mutation landscapes ranges from 0.751 (at 1 Mb) to 0.894 (at 50 kb, **Supplemental Table S4**). The recombination landscape is also well recovered, with correlation coefficients ranging from 0.830 (at 50 kb) to 0.963 (at 1 Mb, **Supplemental Table S5**). We postulate that iSMC’s power to distinguish between the signal that θ and τ leave on sequence data stems exactly from the difference in scale at which they vary along the simulated chromosomes. Although linked selection can increase the correlation among genealogies around constrained sites (McVean, 2007), in most genomic regions the extent of such effect is still short in comparison to the scale of mutation rate variation, allowing their effects on the distribution of diversity to be teased apart. In summary, although under linked selection the accuracy of inferred mutation maps is lower than under strict neutrality (cf. **Supplemental Table S1**), it remains high enough to validate the robustness of our new model of mutation rate variation.

**Figure 6.**
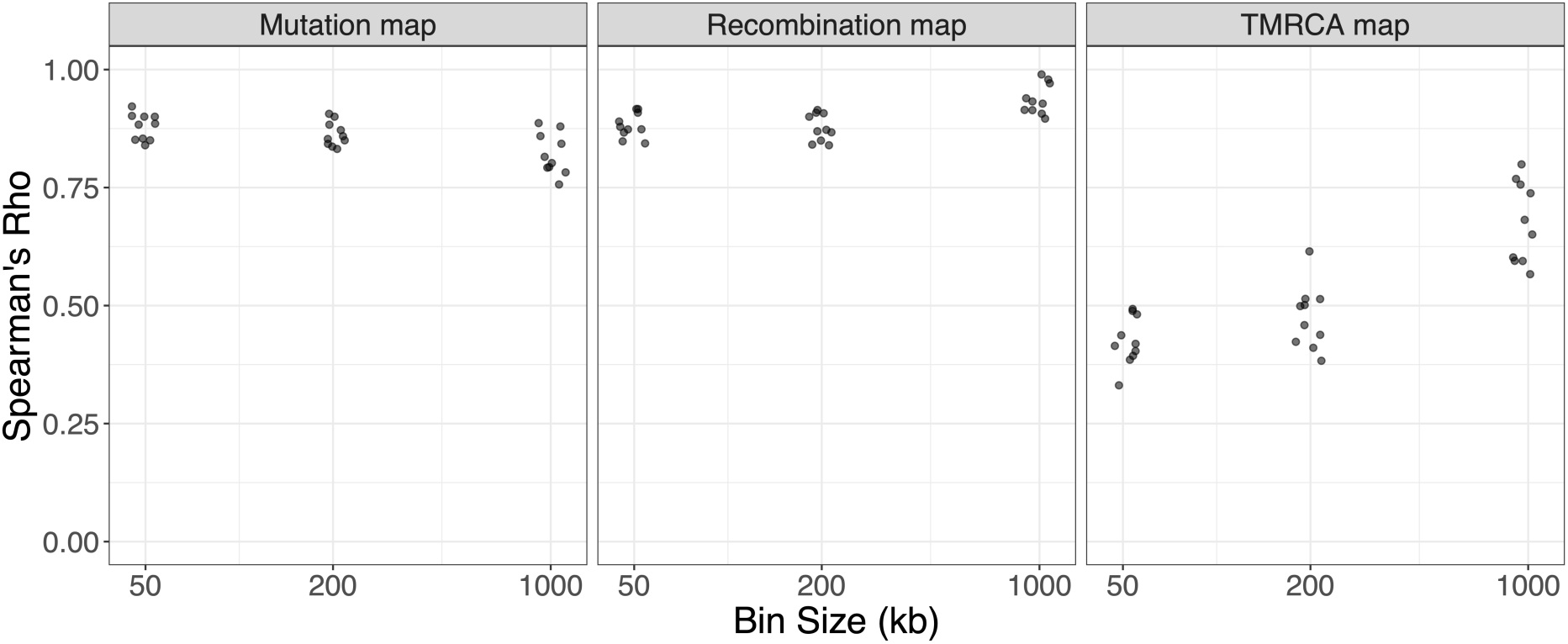
Spearman correlations between inferred and simulated landscapes under background selection. All p-values are smaller than 1e-4.

Given the high accuracy of iSMC in these challenging simulations, one would expect hefty resemblance in the linear models when using the inferred versus the true, noise-free landscapes. This is indeed what we found at all scales (middle and right panels in **Figure 4B**), suggesting that residual biases in iSMC due to linked selection do not carry over to the regression analyses noticeably. Moreover, there is a closer agreement of real *D. melanogaster* data with neutral simulations than with simulations of background selection. This is probably a consequence of the unrealistically strong background selection in the simulations, where the distribution of fitness effects we used has a high density of weakly deleterious mutations which segregate longer in the population, leading to a more localized and pronounced distortion of genealogies (Zeng & Charlesworth, 2011). This would also explain why the R^2^ attributed to θ and τ respectively decrease and increase with window size (**Figure 4B**), a reverse relationship than observed in the real data and in neutral simulations (**Figure 4A**, **Figure 5**). These results are corroborated by the linear coefficients (**Supplemental Table S9**). In the presence of selection, the coefficient of θ decreases with window size and is distinctively closer to the coefficient of τ than under neutrality, a relationship that is reproduced when fitting the linear models with 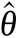 and 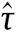. Note also that under intense background selection the coefficient of ρ is generally larger than in the previous scenarios (and the corresponding p-values smaller), mis-matching real *D. melanogaster* data as well. We further inquired into these phenomena visually, by looking into the simulated landscapes, which provided critical insight into the interplay among micro-evolutionary mechanisms shaping diversity (**Figure 7**).

**Figure 7.**
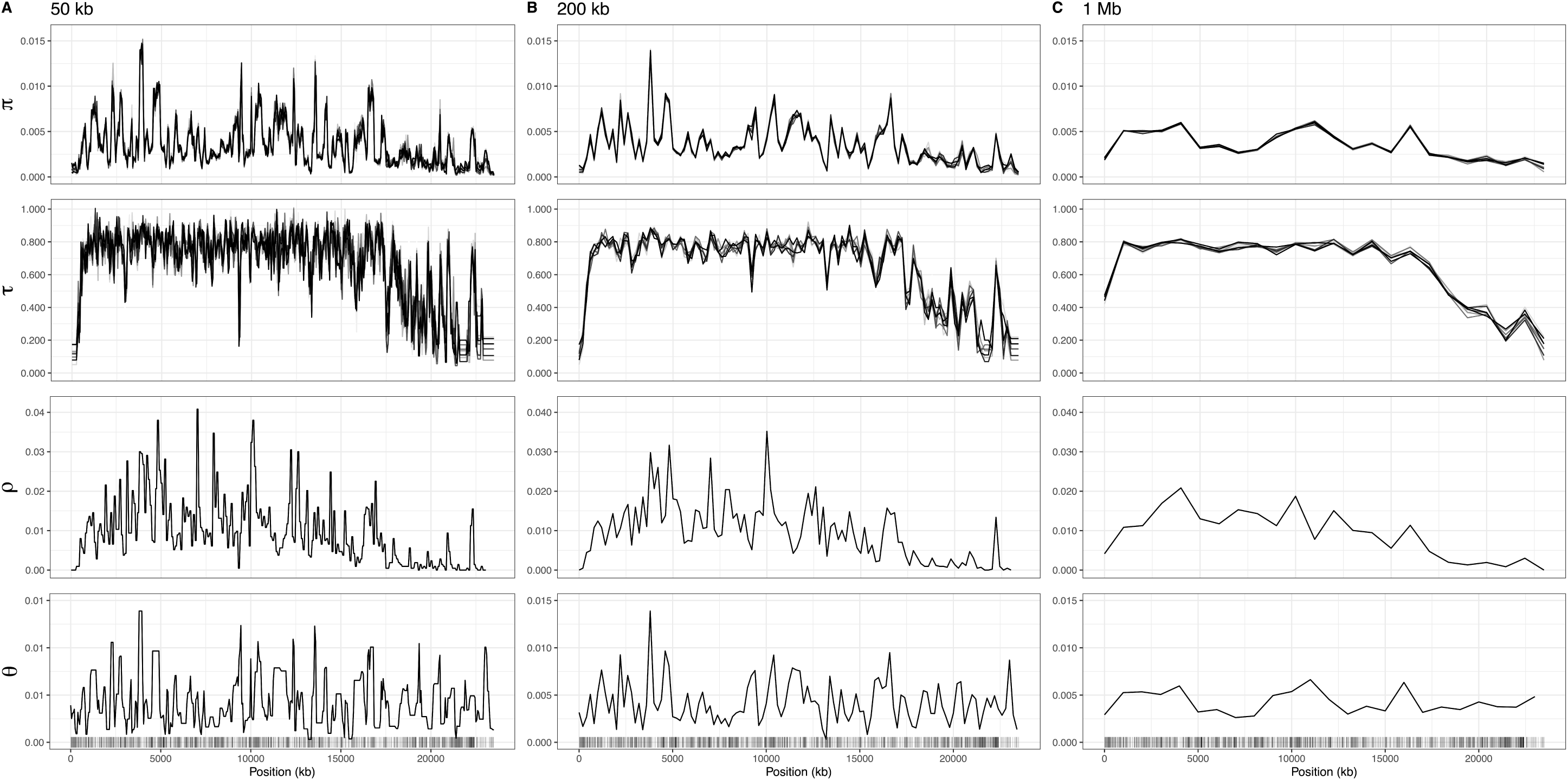
Genomic landscapes simulated under background selection and their effect on the distribution of diversity. From top to bottom: observed nucleotide diversity (lines in shades of grey represent replicates), average TMRCA of the genealogies in units of generations (lines in shades of grey represent replicates), the recombination landscape (shared among replicates) and the mutation landscape (shared among replicates). All landscapes are binned into non-overlapping windows of 50 kb (**A**), 200 kb (**B**) and 1 Mb (**C**). Barcode at the bottom represents the density of sites under negative selection. These data were extracted straight from the simulations and are, therefore, free of estimation noise.

Close inspection shows that only in regions of extremely reduced recombination (the left tip and right tail of the simulated chromosome) does linked selection introduce enough correlation among selected and neutral sites as to influence diversity to a larger degree than mutation rate variation, and this effect seemingly grows with window size. Otherwise, the distribution of π predominantly mirrors that of θ, endorsing our previous results. As a side note, **Figure 7** lays out a rather enticing graphic of our linear models: they seek to represent π (the top row) as a linear combination of τ, ρ, and θ (the other three rows), plus the interaction τ:θ. From this perspective, it becomes apparent that the mutation landscape contributes the most to variation in diversity along the chromosome, even in such a conservative scenario where linked selection is artificially strong. Taken together, the results from these simulations provide compelling evidence that the high R^2^ attributed to θ in *D. melanogaster* is a solid finding. First, it cannot be explained either by the increased noise in our inference of τ compared to θ (**Figure 6**), or by potential absorption of linked selection effects into 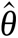, since in both of these cases we would not expect the close correspondence between OLS results fitted with inferred versus true maps (**Figure 4b**). Second, raw simulated data clearly demonstrate that the effect of linked selection can be overwhelmed by mutation rate variation (**Figure 7**). We conclude that the modeling framework illustrated in **Figures 1** and **3** satisfactorily captures the essence of the genome-wide determinants of nucleotide diversity, and is likewise adequate to the study of *D. melanogaster*.

## Discussion

The presence of mutation rate variation along the genome has been recognized for many years (some of the evidence in mammals reviewed over a decade ago by Hodgkinson & Eyre-Walker (2011)), although its implications have been largely overlooked by the population genetics literature. The contributions of the present work are not to simply recapitulate this phenomenon in *D. melanogaster*, but mainly to (1) present a novel statistical method that can infer such variation using population genetic data and (2) use this method to show that the mutation landscape has a lasting effect on nucleotide diversity patterns that can be quantitatively larger than that of natural selection. This awareness is long overdue, as the relative strengths of selection and drift in shaping genome-wide diversity have been debated for several decades (reviewed in Hey (1999); and, more recently, Kern & Hahn (2018) and Jensen et al. (2019)), with the influence of local mutation rate only recently brought to light (Castellano et al., 2018; Harpak et al., 2016; Smith et al., 2018). We were able to employ our extended iSMC model to jointly infer mutation, recombination and TMRCA landscapes and to use causal inference to estimate their impact on π along the genome. Our analyses revealed that these combined landscapes explain >99% of the distribution of diversity along the *D. melanogaster* genome; when looking into the detailed patterns, we found the footprints of linked selection, but the major driver of genome-wide diversity in this species seems to be the mutation landscape. Importantly, this conclusion holds whether we base the discussion on estimates of linear coefficients or on the proportion of variance explained.

These results do not imply that linked selection cannot extend beyond the 18.3% of the *D. melanogaster* genome that is exonic (Alexander et al., 2010), but rather that variation in the mutation rate is strong enough to contribute relatively more to the variation in π, in the genomic scales here employed (**Figure 4**). Our findings, however, sharply contrast with an estimate by Comeron that up to 70% of the distribution of diversity in *D. melanogaster* can be explained solely by background selection at the 100 kb scale (Comeron, 2014), where the author further argued that many regions of increased diversity may be experiencing balancing selection. Instead, we propose that mutation rate variation is responsible for most of these effects. We believe that such discrepancy can be mainly attributed to Comeron’s 70% figure deriving from the (rank) correlation between π and B-value maps alone, without including other causal factors (like drift and local μ). The B-value represents the expected reduction of diversity due to selection against linked deleterious mutations (Charlesworth, 2013; Matheson & Masel, 2022; McVicker et al., 2009). This is equivalent to a scaling of the expected TMRCA between two (uniformly) random samples, which in our model is captured by 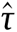. Indeed we find that despite 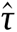 explaining little variance in diversity in the multiple regression setting, the simple correlations between 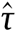 and π are of the same order as found by Comeron (Spearman’s rho = 0.70, p-value < 2.2e-16 at the 50 kb scale; Spearman’s rho = 0.66, p-value < 2.2e-16 at the 200 kb scale; Spearman’s rho = 0.86, p-value < 2.2e-16 at the 1 Mb scale, cf. Table 1 in (Comeron, 2014)). The central point being that the linear models were able to reliably pinpoint θ as the main driver of π just because its effect was jointly estimated with those of τ and ρ. Taking a step back, it is also conceivable that selection is not only manifested as distortions in the distribution of genealogies, but also biases our estimate of the mutation landscape. We note, however, that there actually seems to be a small overestimation of the importance of the TMRCA in our results (compare values obtained with true vs inferred landscapes in **Figure 4** and **Supplemental Tables S2, S9**), which goes in the opposite direction to the presumed bias under linked selection. In this way our results appear to be conservative with respect to the discussions we submitted throughout this article. On top of that, based on the high similarity between real *D. melanogaster* data and our neutral simulations (**Figure 4A**) as well as on iSMC’s robustness to the presence of linked selection (**Figure 2B**, **Figure 4B**, **Figure 6**), we argue that a bias induced by linked selection would likely be insufficient to overturn our conclusion of a major impact of mutation rate variation on the distribution of diversity.

We also note that selection should have a stronger impact on π when binning is performed at smaller genomic scales (≤10 kb, e.g., Figure 4 in Hudson & Kaplan (1995)), which we have not explored because of increased genealogical and mutational variance at such small window sizes. Besides Comeron, Elyashiv et al. (2016) also used patterns of nucleotide variation to fit models of linked selection along the fruit fly genome. Using substitution rates at synonymous sites as a proxy for local mutation rates, they employed their selection estimates to predict genome-wide diversity in windows from 1 kb to 1 Mb. Their models predict 44% (100 kb) and 76% (1 Mb) of the distribution of scaled nucleotide diversity in *D. melanogaster*. However, owing to the scaling that removes the effect of mutation rate variation, the percentages in the Elyashiv et. al study represent the part of variance explained by linked selection once the effect of mutation rate variation has been discarded (see also Murphy et al. (2022) for a similar and improved model). As such, the R^2^ values they report quantify the goodness-of-fit of different models of selection (e.g., background selection alone vs background selection + selective sweeps) instead of the actual importance of linked selection to π, and are, therefore, not directly comparable with our estimates. Still, we note that the remaining variance in their models may be due to mutation rate variation not grasped by synonymous divergence – an imperfect proxy for μ, either because of selection on codon usage or because the mutation landscape has evolved since the divergence of the two species. (Along the same vein, the correlation between our 50 kb mutation maps and genome-wide divergence between *D. melanogaster* and *D. yakuba* is only moderate, Spearman’s rho = 0.197, p-value = 3e-09.) The differences between our approaches to capture linked selection are also worth discussing. While Elyashiv et al. (2016) relied on elaborate models of selection that embody strong assumptions, we leaned on the spatial distribution of τ, similarly to Palamara et al. (2018). This heuristic renders our approach more parsimonious (11 parameters compared to 36 in the Elyashiv et. al model) and less susceptible to mis-specifications of the selection model, which could be commonplace (e.g., the presence of epistasis and/or fluctuating fitness effects over time). Developing an explicit model of spatial variation in *N*_e_ into the iSMC framework is desirable but presents considerable obstacles, and is therefore left as a future perspective.

Our results provide evidence that similarly to humans (Harpak et al., 2016; Smith et al., 2018), the mutation landscape is a crucial determinant of the distribution of diversity in *D. melanogaster*. The simulation study (**Figure 5**, **Figure 7**) further suggests that in many evolutionary scenarios the mutation landscape will remain the most relevant factor shaping π along the genome, depending notably on the window size used in the analysis. Future work using integrative models like the ones we introduced here (**Figure 1**, **Figure 3**) and applied to species with distinct genomic features and life-history traits will help elucidate how often – and by how much – the mutation landscape stands out as the main driver of nucleotide diversity.

We emphasize that we have not directly argued in favor of either genetic drift or natural selection in the classic population genetics debate, but instead we have highlighted the importance of a third element – the mutation landscape – in shaping genome-wide diversity. Nevertheless, the mutation landscape should play a role in the dynamics of natural selection by modulating the rate at which variation is input into genes (and other functionally important elements) depending on their position in the genome. Consequently, levels of selective interference, genetic load and rates of adaptation should vary accordingly (Castellano et al., 2016). In *D. melanogaster*, our inferred mutation landscape varies ∼10fold between minimum and maximum values at the 50 kb scale, meaning that the impact of mutation rate variation on selective processes can be substantial. This opens intriguing lines of inquiry. For example, under what conditions can the shape of the mutation landscape itself be selected for? It has been shown that modifiers of the global mutation rate are under selection to reduce genetic load (Lynch, 2008; Lynch et al., 2016). It remains to be seen whether the position of genes or genomic features correlated with local μ (e.g., replication timing (Francioli et al., 2015)) can likewise be optimized (Martincorena & Luscombe, 2013). After all, population genetics theory predicts that at equilibrium the reduction in mean fitness of the population due to recurrent mutations is equal to the sum of mutation rates among sites where they hit deleteriously and actually independent of their selective effects (Haldane, 1937). Curiously, during the editing of this manuscript, the first evidence of an adaptive mutation landscape was reported in *Arabidopsis thaliana*, with coding regions experiencing fewer *de novo* mutations than the rest of the genome, and essential genes even less so (Monroe et al., 2022). This suggests that local mutation rates have been themselves under selective pressure to reduce genetic load in at least one model system, and indicates that perhaps an even smaller fraction of the depletion of nucleotide diversity near genes can be directly attributed to linked selection than previously inferred. In *D. melanogaster*, we failed to find a relationship between local mutation rate and selective constraint (recall that the correlation test between 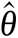 and the proportion of exonic sites yielded Spearman’s rho = – 0.037 with p-value = 0.19, at the 50 kb scale); however, this could also be due to lack of power in the test because of the relatively large window size we used, combined with *Drosophila*’s high gene density. At any rate, much more effort is needed to explore the causes and consequences of mutation rate variation across the tree of life. As a starting point, we can ask how conserved the mutation landscape is in closely-related species (or, equivalently, how fluid is its evolution within populations). Analogous work on the recombination landscape has revealed overall fast evolution of “hotspots” in mammals and has helped uncover the molecular architecture responsible for the placement of double-strand breaks (Berg et al., 2011; Jabbari et al., 2019). Moreover, adaptive dynamics have been evoked to explain the differences in the recombination landscape between populations of *D. pseudoobscura*, (Samuk et al., 2020). It will be interesting to test whether mutation events follow similar patterns, now that the impact of various sequence motifs on local μ is being more thoroughly investigated (DeWitt et al., 2021; Kim et al., 2021; Oman et al., 2022). Unraveling the factors that shape the mutation landscape at different genomic scales will likely provide important insight. For example, can the large-scale variation in mutation rates that we found in *D. melanogaster* be partially explained by aggregation of short (differentially mutable) sequence motifs, or is it driven by independent genomic features? As the molecular underpinnings of adaptive mutation landscapes become elucidated (e.g., what kind of proteins, sequence motifs and epigenetic markers are involved in increasing replication fidelity in functionally constrained regions and eventually decreasing it where polymorphism would tend to be beneficial) we will gain a better understanding of how flexible such phenotype is and how prevalent it is expected to be in different phylogenetic groups. It is plausible that modifiers of the mutation landscape may be successfully optimized, at least in species with high enough *N*_e_ for such second-order effects to be seen by selection (Lynch, 2010; Martincorena & Luscombe, 2013; Sung et al., 2012). Recent work notably highlighted the importance of epigenetic factors in shaping the mutation landscape and started to shed light on its evolutionary consequences (Habig et al., 2021; Möller et al., 2021). The variety of molecular agents recruited to tweak the mutation landscape and create pockets of decreased or increased *de novo* mutation rates can be plenty, and it only outlines the complexity of evolutionary biology.

In hindsight, it is perhaps not surprising that mutation rate variation has a profound impact on nucleotide diversity. Mutations are, after all, the “stuff of evolution” (Nei, 2013), and distinct genomic regions displaying differential influx of SNPs must have sharp consequences to the analyses and interpretation of genetic data. This argument is naturally transferred to the ongoing discussion about incorporating complex demography and background selection into the null model of molecular evolution (Comeron, 2017, 2014; Johri et al., 2020), which is motivated by the goal of providing more sensible expectations for rigorously testing alternative scenarios. Our results suggest that a more realistic null model should also include variation in the mutation rate along the genome. By doing so, genome-wide scans (e.g., looking for regions with reduced diversity summary statistics as candidates for selective sweeps) may become less susceptible to both false negatives (in regions of high mutation rate) and false positives (in regions of low mutation rate), paving the way to more robust inference (Booker et al., 2017; Haasl & Payseur, 2016; Venkat et al., 2018).

## Acknowledgments

The authors thank Aaron Ragsdale, Chris Kyriazis, Guillaume Achaz, Guy Sella, Kai Zeng, Laurent Duret, Josep Comeron, Kirk Lohmueller and Pier Palamara for discussions about this work; Ana Filipa Moutinho for providing organized data on *Drosophila*.

## Funding

JYD acknowledges funding from the Max Planck Society. This work was supported by a grant from the German Research Foundation (Deutsche Forschungsgemeinschaft) attributed to JYD, within the priority program (SPP) 1590 “probabilistic structures in evolution”.

## Conflicts of Interest Statement

The authors declare they have no conflict of interest relating to the content of this article.

## Data and Software Availability

The iSMC software package and source code is freely available at https://github.com/gvbarroso/iSMC. The DOI for this repository is 10.5281/zenodo.7826556. Scripts used to generate our results can be found in https://github.com/gvbarroso/ismc_dm_analyses. The DOI for this repository is 10.5281/zenodo.7826575. Data required to reproduce our results are deposited in FigShare under the DOI 10.6084/m9.figshare.13164320. The second repository contains the script ‘dm_analyses.Rmd’. Combined with the data on FigShare, this script generates a PDF document named ‘dm_analyses.pdf’ which contains the results and diagnostics of all linear models and correlations reported in this article.

## Notes

https://figshare.com/articles/dataset/Quantifying_the_determinants_of_the_genome-wide_diversity_in_Drosophila_using_iSMC/13164320

### Competing Interest Statement

The authors have declared no competing interest.

### Summary of Updates

Prepares article in the PCI format

