## Supplemental_File_1 for "The landscape of nucleotide diversity in *Drosophila melanogaster* is shaped by mutation rate variation"

**Supplemental Table S1.** Spearman's correlations between inferred and simulated mutation maps. Neutral scenario (coalescent simulations). All p-values < 1e-4.

| Replicate/Scale | 50 kb | 200 kb | 1 Mb |
|---|---|---|---|
| 1 | 0.978 | 0.985 | 0.979 |
| 2 | 0.985 | 0.989 | 0.975 |
| 3 | 0.978 | 0.983 | 0.981 |
| 4 | 0.978 | 0.982 | 0.982 |
| 5 | 0.982 | 0.987 | 0.983 |
| 6 | 0.984 | 0.987 | 0.981 |
| 7 | 0.98 | 0.987 | 0.975 |
| 8 | 0.981 | 0.988 | 0.982 |
| 9 | 0.983 | 0.987 | 0.981 |
| 10 | 0.984 | 0.989 | 0.984 |

**Supplemental Table S2. Estimates from linear models fitted to the distribution of nucleotide diversity along genomes simulated under the neutral coalescent.** Vertical panels show results according to genomic window size whereas horizontal panels show results according to the origin of the landscapes used to fit the linear models (either true or iSMC-inferred). Explanatory variables were standardised prior to model fitting. Since all 10 replicates yielded highly consistent estimates, we only show results from the first replicate. OLS = Ordinary Least Squares, GLS = Generalised Least Squares, VIF = Variance Inflation Factor.

| | | | 50 kb | | | 200 kb | | | 1 Mb | | |
|---|---|---|---|---|---|---|---|---|---|---|---|
| **Map Source** | **Type** | **Variable** | **Coefficient** | **p-value** | **VIF** | **Coefficient** | **p-value** | **VIF** | **Coefficient** | **p-value** | **VIF** |
| **Simulated** | **OLS** | $\theta$ | 0.0100 | <2.2e-16 | 1.01 | 0.0097 | <2.2e-16 | 1.01 | 0.0064 | <2.2e-16 | 1.25 |
| | | $\tau$ | 0.0017 | <2.2e-16 | 1.00 | 0.0009 | <2.2e-16 | 1.00 | 0.0004 | 0.0000 | 1.12 |
| | | $\rho$ | 0.0000 | 0.0266 | 1.01 | 0.0005 | 0.0890 | 1.02 | 0.0000 | 0.9700 | 1.04 |
| | | $\theta:\tau$ | 0.0089 | <2.2e-16 | 1.01 | 0.0004 | <2.2e-16 | 1.01 | 0.0004 | 7e-02 | 1.37 |
| **Inferred** | **OLS** | $\theta$ | 0.0100 | <2.2e-16 | 1.02 | 0.0093 | <2.2e-16 | 1.02 | 0.0063 | <2.2e-16 | 1.08 |
| | | $\tau$ | 0.0028 | <2.2e-16 | 1.08 | 0.0022 | <2.2e-16 | 1.28 | 0.0009 | <2.2e-16 | 1.93 |
| | | $\rho$ | 0.0000 | 0.8000 | 1.01 | 0.0000 | 0.8230 | 1.07 | 0.0000 | 0.9930 | 1.28 |
| | | $\theta:\tau$ | 0.0014 | <2.2e-16 | 1.06 | 0.0011 | <2.2e-16 | 1.18 | 0.0003 | 0.113 | 1.73 |
| | **GLS** | $\theta$ | 0.0104 | <1e-4 | 1.03 | 0.0093 | <1e-4 | 1.02 | 0.0063 | <1e-4 | 1.09 |
| | | $\tau$ | 0.0028 | <1e-4 | 1.05 | 0.0022 | <1e-4 | 1.25 | 0.0009 | <1e-4 | 1.90 |
| | | $\rho$ | 0.0000 | 0.8355 | 1.00 | 0.0000 | 0.7295 | 1.07 | 0.0000 | 0.9982 | 1.27 |
| | | $\theta:\tau$ | 0.0014 | <1e-4 | 1.04 | 0.0011 | <1e-4 | 1.68 | 0.0003 | 0.1041 | 1.72 |

**Supplemental Table S3. Spearman's correlations between inferred and simulated TMRCA maps.** Background selection scenario (forward simulations with SLiM), all p-values < 1e-4.

| Replicate/Scale | 50 kb | 200 kb | 1 Mb |
|---|---|---|---|
| 1 | 0.385 | 0.442 | 0.666 |
| 2 | 0.421 | 0.403 | 0.610 |
| 3 | 0.449 | 0.461 | 0.783 |
| 4 | 0.343 | 0.423 | 0.586 |
| 5 | 0.400 | 0.426 | 0.641 |
| 6 | 0.367 | 0.466 | 0.700 |
| 7 | 0.424 | 0.485 | 0.614 |
| 8 | 0.443 | 0.463 | 0.787 |
| 9 | 0.465 | 0.569 | 0.745 |
| 10 | 0.408 | 0.545 | 0.760 |

**Supplemental Table S4. Spearman's correlations between inferred and simulated mutation maps.** Background selection scenario (forward simulations with SLiM), all p-values < 1e-4.

| Replicate/Scale | 50 kb | 200 kb | 1 Mb |
|---|---|---|---|
| 1 | 0.894 | 0.873 | 0.813 |
| 2 | 0.851 | 0.841 | 0.840 |
| 3 | 0.872 | 0.858 | 0.764 |
| 4 | 0.882 | 0.875 | 0.777 |
| 5 | 0.870 | 0.863 | 0.783 |
| 6 | 0.863 | 0.853 | 0.801 |
| 7 | 0.873 | 0.863 | 0.751 |
| 8 | 0.871 | 0.859 | 0.842 |
| 9 | 0.886 | 0.880 | 0.803 |
| 10 | 0.883 | 0.869 | 0.810 |

**Supplemental Table S5. Spearman's correlations between inferred and simulated recombination maps.** Background selection scenario (forward simulations with SLiM), all p-values < 1e-4.

| Replicate/Scale | 50 kb | 200 kb | 1 Mb |
|---|---|---|---|
| 1 | 0.870 | 0.879 | 0.938 |
| 2 | 0.879 | 0.903 | 0.963 |
| 3 | 0.868 | 0.894 | 0.935 |
| 4 | 0.870 | 0.898 | 0.952 |
| 5 | 0.881 | 0.896 | 0.916 |
| 6 | 0.830 | 0.863 | 0.957 |
| 7 | 0.865 | 0.879 | 0.942 |
| 8 | 0.871 | 0.914 | 0.943 |
| 9 | 0.872 | 0.888 | 0.940 |
| 10 | 0.851 | 0.873 | 0.928 |

**Supplemental Table S6.** Generalised Least Squares models fit to *Drosophila melanogaster* autosomes using 50 kb scale maps (* denotes statistical significance at 5% level after Bonferroni correction)

| Chr | Variable | Coefficient | p-value | Sig? |
|---|---|---|---|---|
| | θ | 0.0023 | <1e-4 | * |
| 2L | τ | 0.0008 | <1e-4 | * |
| | ρ | 0.0000 | 0.2250 | NS |
| | θ:τ | 0.0002 | <1e-4 | * |
| | θ | 0.0023 | <1e-4 | * |
| 2R | τ | 0.0007 | <1e-4 | * |
| | ρ | 0.0000 | 0.9078 | NS |
| | θ:τ | 0.0001 | <1e-4 | * |
| | θ | 0.0029 | <1e-4 | * |
| 3L | τ | 0.0012 | <1e-4 | * |
| | ρ | 0.0000 | <1e-4 | |
| | θ:τ | 0.0003 | <1e-4 | * |
| | θ | 0.0026 | <1e-4 | * |
| 3R | τ | 0.0011 | <1e-4 | * |
| | ρ | 0.0000 | 0.1449 | NS |
| | θ:τ | 0.0003 | <1e-4 | * |

**Supplemental Table S7.** Generalised Least Squares models fit to *Drosophila melanogaster* autosomes using 200 kb scale maps (* denotes statistical significance at 5% level after Bonferroni correction)

| Chr | Variable | Coefficient | p-value | Sig? |
|---|---|---|---|---|
| 2L | θ | 0.0020 | <1e-4 | * |
| | τ | 0.0006 | <1e-4 | * |
| | ρ | 0.0000 | 0.2807 | NS |
| | θ:τ | 0.0001 | <1e-4 | * |
| 2R | θ | 0.0025 | <1e-4 | * |
| | τ | 0.0004 | <1e-4 | * |
| | ρ | 0.0000 | 0.3516 | NS |
| | θ:τ | 0.0001 | <1e-4 | * |
| 3L | θ | 0.0028 | <1e-4 | * |
| | τ | 0.0007 | <1e-4 | * |
| | ρ | 0.0000 | 0.0037 | * |
| | θ:τ | 0.0002 | <1e-4 | * |
| 3R | θ | 0.0026 | <1e-4 | * |
| | τ | 0.0010 | <1e-4 | * |
| | ρ | 0.0000 | 0.0211 | NS |
| | θ:τ | 0.0002 | <1e-4 | * |

**Supplemental Table S8.** Generalised Least Squares models fit to *Drosophila melanogaster* autosomes using 1 Mb scale maps (* denotes statistical significance at 5% level after Bonferroni correction)

| Chr | Variable | Coefficient | p-value | Sig? |
|---|---|---|---|---|
| | θ | 0.0015 | <1e-4 | * |
| 2L | τ | 0.0004 | <1e-4 | * |
| | ρ | 0.0000 | 0.8532 | NS |
| | θ:τ | 0.0001 | <1e-4 | * |
| | θ | 0.0019 | <1e-4 | * |
| 2R | τ | 0.0002 | <1e-4 | * |
| | ρ | 0.0000 | 0.3479 | NS |
| | θ:τ | 0.0000 | 0.3277 | NS |
| | θ | 0.0026 | <1e-4 | * |
| 3L | τ | 0.0005 | <1e-4 | * |
| | ρ | 0.0000 | 0.0913 | NS |
| | θ:τ | 0.0001 | 0.0298 | NS |
| | θ | 0.0026 | 0.0000 | * |
| 3R | τ | 0.0010 | 0.0000 | * |
| | ρ | 0.0000 | 0.2173 | NS |
| | θ:τ | 0.0002 | 0.0010 | * |

**Supplemental Table S9. Estimates from linear models fitted to the distribution of nucleotide diversity along genomes simulated under background selection.** Vertical panels show results according to genomic window size whereas horizontal panels show results according to the origin of the landscapes used to fit the linear models (either true or iSMC-inferred). Explanatory variables were standardised prior to model fitting. Since all 10 replicates yielded highly consistent estimates, we only show results from the first replicate. OLS = Ordinary Least Squares, GLS = Generalised Least Squares, VIF = Variance Inflation Factor.

| | | | **50 kb** | | | **200 kb** | | | **1 Mb** | | |
|---|---|---|---|---|---|---|---|---|---|---|---|
| **Map Source** | **Type** | **Variable** | **Coefficient** | **p-value** | **VIF** | **Coefficient** | **p-value** | **VIF** | **Coefficient** | **p-value** | **VIF** |
| **Simulated** | **OLS** | θ | 0.0020 | <2.2e-16 | 1.11 | 0.0016 | <2.2e-16 | 1.14 | 0.0008 | 2.60e-13 | 1.80 |
| | | τ | 0.0010 | <2.2e-16 | 1.70 | 0.0010 | <2.2e-16 | 2.04 | 0.0009 | 6.40e-12 | 3.24 |
| | | ρ | 0.0001 | 0.0007 | 1.70 | 0.0001 | 0.0193 | 2.09 | 0.0002 | 0.0212 | 3.45 |
| | | θ:τ | 0.0007 | <2.2e-16 | 1.11 | 0.0006 | <2.2e-16 | 1.14 | 0.0004 | 2.40e-05 | 1.65 |
| **Inferred** | **OLS** | θ | 0.0020 | <2.2e-16 | 1.24 | 0.0017 | <2.2e-16 | 1.02 | 0.0010 | 1.23e-10 | 7.11 |
| | | τ | 0.0009 | <2.2e-16 | 1.23 | 0.0008 | <2.2e-16 | 1.28 | 0.0007 | 4.69e-13 | 2.11 |
| | | ρ | 0.0001 | <2.2e-16 | 1.34 | 0.00008 | 2.4e-5 | 1.07 | 0.0001 | 0.0301 | 2.97 |
| | | θ:τ | 0.0006 | <2.2e-16 | 1.13 | 0.0005 | <2.2e-16 | 1.18 | 0.0004 | 0.0419 | 5.60 |
| | **GLS** | θ | 0.0021 | <1e-4 | 1.16 | 0.0017 | <1e-4 | 1.02 | 0.0018 | <1e-4 | 1.17 |
| | | τ | 0.0009 | <1e-4 | 1.15 | 0.0008 | <1e-4 | 1.25 | 0.0007 | <1e-4 | 1.23 |
| | | ρ | 0.0001 | 0.0002 | 1.11 | 0.00009 | <1e-4 | 1.07 | 0.00005 | 0.0079 | 1.33 |
| | | θ:τ | 0.0006 | <1e-4 | 1.14 | 0.0004 | <1e-4 | 1.68 | 0.00036 | <1e-4 | 1.06 |

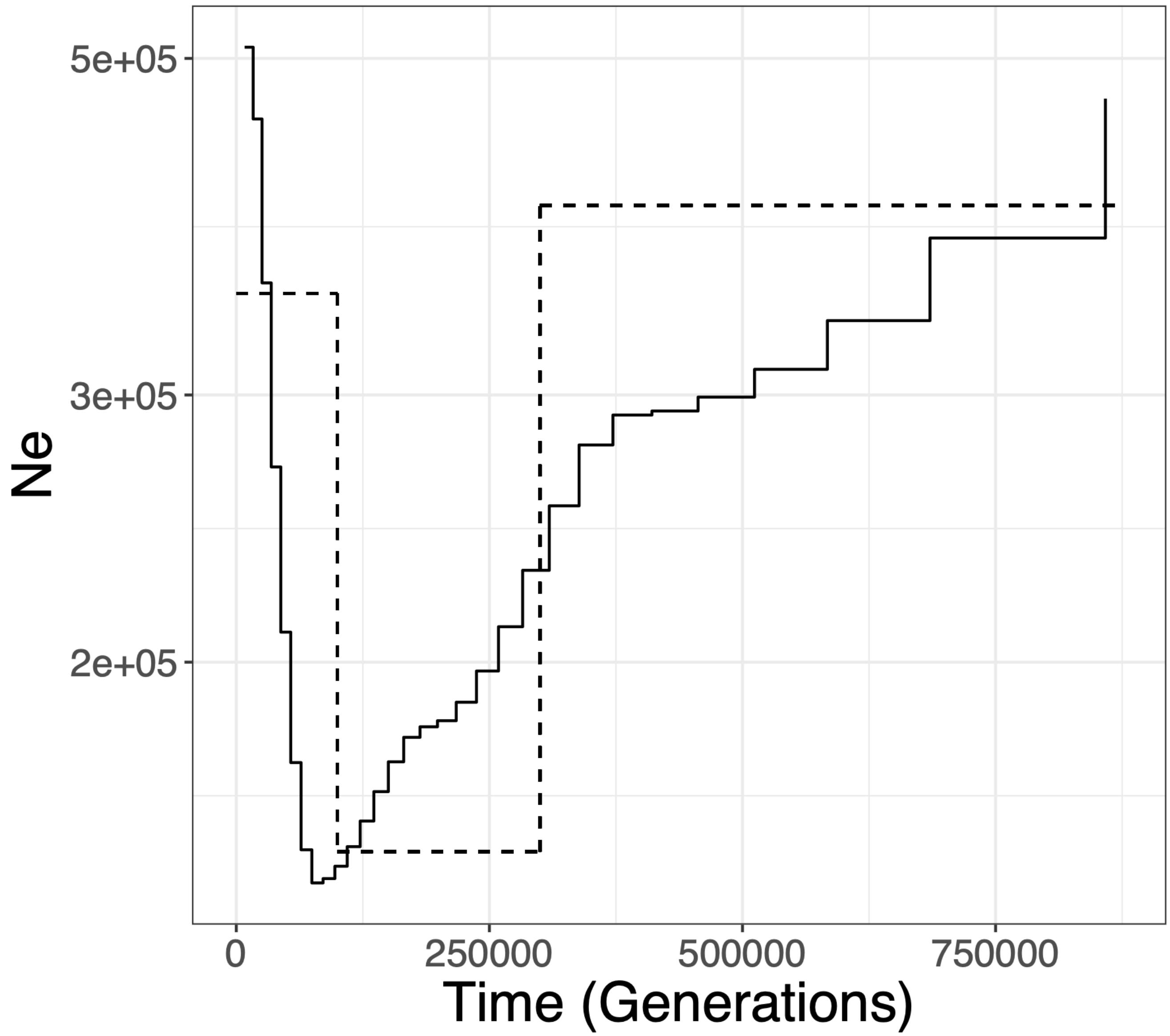


**Supplemental Figure S1. Demographic history of the Zambia population of *Drosophila melanogaster*.** Solid line shows population size trajectory inferred using iSMC, dashed line shows smoothed scheme used to perform coalescent simulations.
